# Assembly and Conformational Dynamics of Insect Odorant Receptors

**DOI:** 10.64898/2026.08.16.745119

**Authors:** Yining Jiang, Jiawei Zhao, Zhaokun Wang, Josefina del Mármol, Simon Scheuring

## Abstract

Insects rely on odorant receptors (ORs), tetrameric ligand-gated ion channels, to detect and discriminate a wide range of chemical cues. Within the insect OR family, composed of hundreds of OR subtypes, a conserved subunit termed odorant receptor co-receptor (Orco) is present in insect olfactory neurons. While Orco and some ORs can assemble into homo-tetramers, odor-activated channels are hetero-tetramers comprising Orco and an odor-specific OR. Cryogenic electron microscopy (cryo-EM) structures from heterologous expression systems revealed that Orco and OR assembled with a 3:1 (Orco:OR) stoichiometry. In these complexes, the OR subunit is solely responsible for ligand-recognition, whereas the Orco subunits form the majority of the ion conduction pathway and are key for assembly and membrane trafficking. Because Orco and OR can assemble in diverse stoichiometries, of which Orco_(4)_ and OR_(4)_ homo-tetramers, as well as Orco_(3)_/OR_(1)_ hetero-tetramers, have been experimentally documented, we sought to investigate assembly and conformational dynamics of insect Orco/OR complexes. Here, using high-speed atomic force microscopy (HS-AFM), unique in providing simultaneous structural and temporal resolution, we find that Orco/OR complexes can reversibly transition into stable non-tetrameric oligomeric states, such as trimers and pentamers, through membrane-diffusive oligomer exchange. Furthermore, while all cryo-EM structures showed a fourfold symmetric or pseudo-symmetric conformation, we observed that tetrameric assemblies frequently adopt a symmetry-broken ‘pinched’ configuration, in addition to the canonical symmetric ‘square’ arrangement. Notably, ligand-binding modulates the ‘pinching’ kinetics in Orco/OR hetero-tetramers. Overall, these findings reveal key aspects of assembly and conformational dynamics of the insect OR family.

## Introduction

Insects are remarkably capable of odor discernment, allowing them to effectively navigate, communicate, and locate food sources based on chemical cues^1^. Each insect olfactory neuron expresses a narrow subset of receptors (from a large family of receptors) that detect a limited set of ligands^2^, and odor discrimination arises from the combinatorial activation patterns across many olfactory neurons^3–5^. This organizational scheme sets the foundation for animals to respond to a diverse chemical world with a finite number of receptors. In contrast to mammalian olfaction, which relies mostly on G protein-coupled receptors (GPCRs), insect odorant receptors (ORs) form ligand-gated ion channels^6–8^. A diverse repertoire of ORs has been identified in insects, ranging from a handful in damselfly to several hundreds in ants^6^.

Insect ORs exhibit striking sequence variation, with only ∼20% amino acids shared across ORs between and within species, which confer them broad ligand diversity^9^. Among the insect OR repertoire, a single component known as the odorant receptor co-receptor (Orco) stands out for its exceptional sequence conservation in nearly all insect species and ubiquitous expression across olfactory neurons^10^. While responding to no known natural odorant, Orco is indispensable for proper odor detection by ORs in most insects. In its absence, ORs fail to assemble and traffic to dendritic membranes, or operate as functional receptors^10,11^. Consequently, Orco mutations lead to severe olfactory defects^12^.

Previous biochemical and structural studies have established that insect ORs assemble as tetramers. In heterologous expression systems, Orco and a few ORs from species lacking Orco in their genome can form homo-tetramers^6,13^. Remarkably, the cryo-EM structure of these homo-tetramers revealed a loosely packed transmembrane domain (TMD), with a cloverleaf-shaped cross-section (**Supplementary Figure 1a,b**)^6,13^: Indeed, protomer-protomer interfaces are only stabilized by a single transmembrane helix pair, leaving substantial inlets between neighboring subunits that are presumably filled by lipids. To maintain tetramer integrity, the four subunits are tethered at the cytosolic side via a densely packed ‘anchor’ domain. In contrast, subunits of canonical tetrameric ion channels, such as voltage-gated potassium (Kv) channels, surround the central pore by a tightly packed TMD with large subunit interface areas^14^.

Functional insect olfactory channels are hetero-oligomers composed of Orco and an odor-specific OR. Cryo-EM of heterologous expressed channels revealed a unique structure with 3:1 (Orco(3)/OR(1)) stoichiometry (**Supplementary Figure 1c-e**)^15–18^. These structures further showed that, despite their sequence divergence, ORs share a conserved TMD-fold and protomer-protomer interface with Orco. Moreover, binding of odorant ligands induced a conformational change in the OR subunit that results in a displacement of the OR pore helix, widening the ion-conductive pathway in an asymmetric manner, leaving the TMD architecture largely unchanged^15,16^. Several recent publications of both conspecific and mixed-species hetero-oligomers also showed the same stoichiometry and overall activation mechanism^17,18^. While overall quite similar to Orco, ORs lack both the cytosolic loops extending from the anchor-domain and extracellular surface protruding moieties of Orco, such as a beta-sheet partially resolved in the Orco_(3)_/OR_(1)_ structure and fully predicted by AlphaFold3^19^ (**Supplementary Figure 1**).

Cryo-EM has provided extensive structural insights into insect ORs, yet the physiological role of the Orco_(3)_/OR_(1)_ stoichiometry is under debate, and their dynamic behavior and gating mechanism remain poorly understood. To address these questions, we employed high-speed atomic force microscopy (HS-AFM^20,21^), uniquely capable of delivering simultaneous structural and temporal resolution, to investigate the assembly and conformational dynamics of insect Orco/OR complexes, including fig wasp *Apocrypta bakeri* Orco homomeric receptors and *Apocrypta bakeri* Orco/ mosquito *Anopheles gambiae* OR28 heteromeric receptors. Strikingly, HS-AFM single-molecule imaging reveals stable non-tetrameric oligomeric states of these receptor complexes which have eluded previous cryo-EM characterization. We also observe dynamic assembly rearrangement events, particularly reversible transitions between oligomeric states. On the level of conformational changes, we find that insect OR assemblies frequently adopt a symmetry-broken ‘pinched’ configuration, not previously described, which deviates from the canonical symmetric ‘square’ arrangement observed in cryo-EM studies. The pinched conformation is associated with an increased likelihood of oligomer assembly plasticity. In Orco/OR hetero-tetramers, ligand-binding modulates ‘pinching’ kinetics and stabilizes the square conformation. Through HS-AFM single-molecule structural biology analysis^22^, complemented by computational structure fitting methods that bridge AFM conformations to molecular structures^23^, we gain deeper insights into pinching dynamics, elucidating the involvement of transient substates and the impact of pinching on the central pore architecture.

Overall, HS-AFM dynamic imaging reveals key aspects of assembly and conformational dynamics of insect ORs, highlighting a rich and previously underappreciated structural variability of these complexes.

## Results

### Membrane reconstitution of insect odorant receptors

To gain dynamic insights into insect ORs, we reconstituted homomeric Orco (**Figure 1a**, left, **Supplementary Figure 1a,b**) and heteromeric Orco/OR28 receptors (**Figure 1a**, right, **Supplementary Figure 1c-e**) into membranes consisting of 1,2-dioleoyl-sn-glycero-3-phosphocholine (DOPC), 1,2-dioleoyl-sn-glycero-3-phospho-L-serine (DOPS), and cholesterol at a ratio of 8:1:1 (w:w:w). Negative-stain EM revealed formation of vesicles of roughly 200-500 nm in diameter (**Figure 1b,c**, left), densely packed with receptors (**Figure 1b,c**, right), as judged by the grainy appearance of the proteo-liposome membrane area (**Figure 1c**, right inset). HS-AFM imaging of the proteo-liposomes confirmed successful dense reconstitution (**Figure 1d,e**, left), as individual OR molecules in the membranes were clearly resolved (**Figure 1d,e**, right). Interestingly, these receptors exhibited rapid lateral mobility and a pronounced tendency to form clusters within the membranes (**Supplementary Movies 1**,**2**). Height analysis of the reconstituted membranes from HS-AFM overviews showed that the membrane bilayer had a thickness of ∼4 nm, in agreement with DOPC-rich membranes^24^, and the receptors had a thickness of ∼6.8 nm, in agreement with the cryo-EM structure^6,15^ (**Figure 1f**). Section profile analysis of the receptors revealed uniform membrane insertion with a protrusion height of ∼0.5 nm from the bilayer surface, in agreement with the receptors’ extracellular surface (**Figure 1g**, see **Figure 1a**).

**Figure 1.**
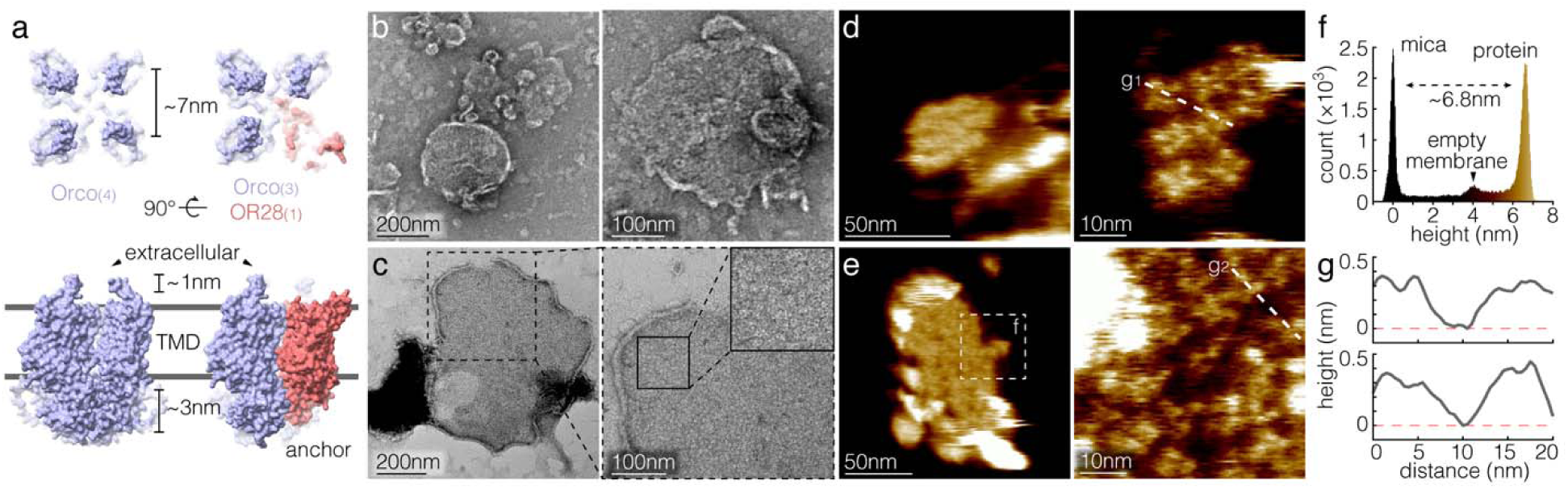
Membrane reconstitution of insect odorant receptors. **(a)** Structural models of homomeric Orco_(4)_ (left) and heteromeric Orco_(3)/_OR28_(1)_ (right) receptors (with full-length extracellular domain, **Supplementary** Figure 1). Top: Top-views of extracellular surfaces. Bottom: Side views. Orco subunits (blue) have an extracellular protrusion, whereas the OR28 subunit (red) lacks such a topographic feature. **(b)** and **(c)** Negative-stain EM micrographs of membrane-reconstituted homomeric Orco (b) and heteromeric Orco/OR28 (c) receptors. Similar results have been obtained from all Orco and Orco/OR28 reconstitution replicates. **(d)** and **(e)** HS-AFM images of the membrane-reconstituted homomeric Orco (d) and heteromeric Orco/OR28 (e) receptors. Left: Overview (**Supplementary Movies 1**), and right: High-resolution images of receptor single molecules. **(f)** Height histogram of the region in (e, dashed outline). **(g)** Height profiles along dashed lines in (d) and (e).

Notably, from the extracellular side, HS-AFM resolved Orco subunits as individual protrusion from the membrane, emerging from the extracellular beta-sheet (**Figure 1a**, blue, **Supplementary Figure 1a,b**). In contrast, OR subunits, due to their lack of a comparable extracellular surface protruding feature (**Figure 1a**, red, **Supplementary Figure 1c-e**), are largely hidden from HS-AFM detection. Therefore, Orco_(4)_ homo-tetramers are cross-shaped and exhibit four visible subunits in HS-AFM (**Figure 1d**, right), while Orco_(3)_/OR28_(1)_ hetero-tetramers display only three visible subunits and one apparent gap (**Figure 1e**, right). Orco subunit protrusions have an inter-protomer distance of ∼5-7 nm in HS-AFM topographies, also consistent with surface representations of the extracellular side of the protein (**Figure 1a**, blue). On the cytosolic side, the anchor-domains of the four subunits tightly bundle together in a unique surface protruding structure (**Supplementary Figure 1**). Thus, the surface protrusion morphology with ∼5-7 nm inter-subunit distance and the subunit protrusion topography height of ∼0.5 nm both agree with the assignment that the tip-exposed surface in HS-AFM experiments is the extracellular side.

### Non-tetrameric assemblies of Orco homomeric receptors

Insect ORs are structurally quite different from other tetrameric ion channels: whilst other channel families have TMDs constituted of densely packed helix bundles that make extended inter-subunit contacts in the membrane, ORs make very little transmembrane helix contacts, and the complex is almost exclusively held together by the cytosolic domain, which was accordingly named anchor-domain (**Figure 2a**). This unique architecture may confer OR oligomer plasticity. Despite the rapid lateral mobility of membrane-reconstituted ORs, HS-AFM fast imaging (10 to 20 frames per second) achieved high-resolution tracking of individual Orco homomeric receptors at the sub-molecular level (**Figure 2b**, **Supplementary Movie 3**). Surprisingly, in addition to molecules that matched the expected appearance of canonical Orco_(4)_ as resolved in cryo-EM studies, we observed the substantial presence of non-tetrameric assemblies (**Figure 2b**, white circles and arrowheads), including pentamers, trimers, dimers and monomers (**Figure 2c**). Statistical analysis of the homomeric Orco oligomer distribution revealed that ∼25-35% of the assemblies were non-tetrameric, with trimers representing the majority (∼60-75% of non-tetramers, ∼15-25% of all oligomers), followed by pentamers (∼20-30% of non-tetramers, ∼5-10% of all oligomers) (**Figure 2d**, Orco WT apo, **Supplementary Table 1**). Therefore, we estimate an equilibrium energy difference of ∼1.2 *k*_B_T between Orco_(4)_ and Orco_(3)_ and ∼2.2 *k*_B_T between Orco_(4)_ and Orco_(5)_ in the experimental conditions.

**Figure 2.**
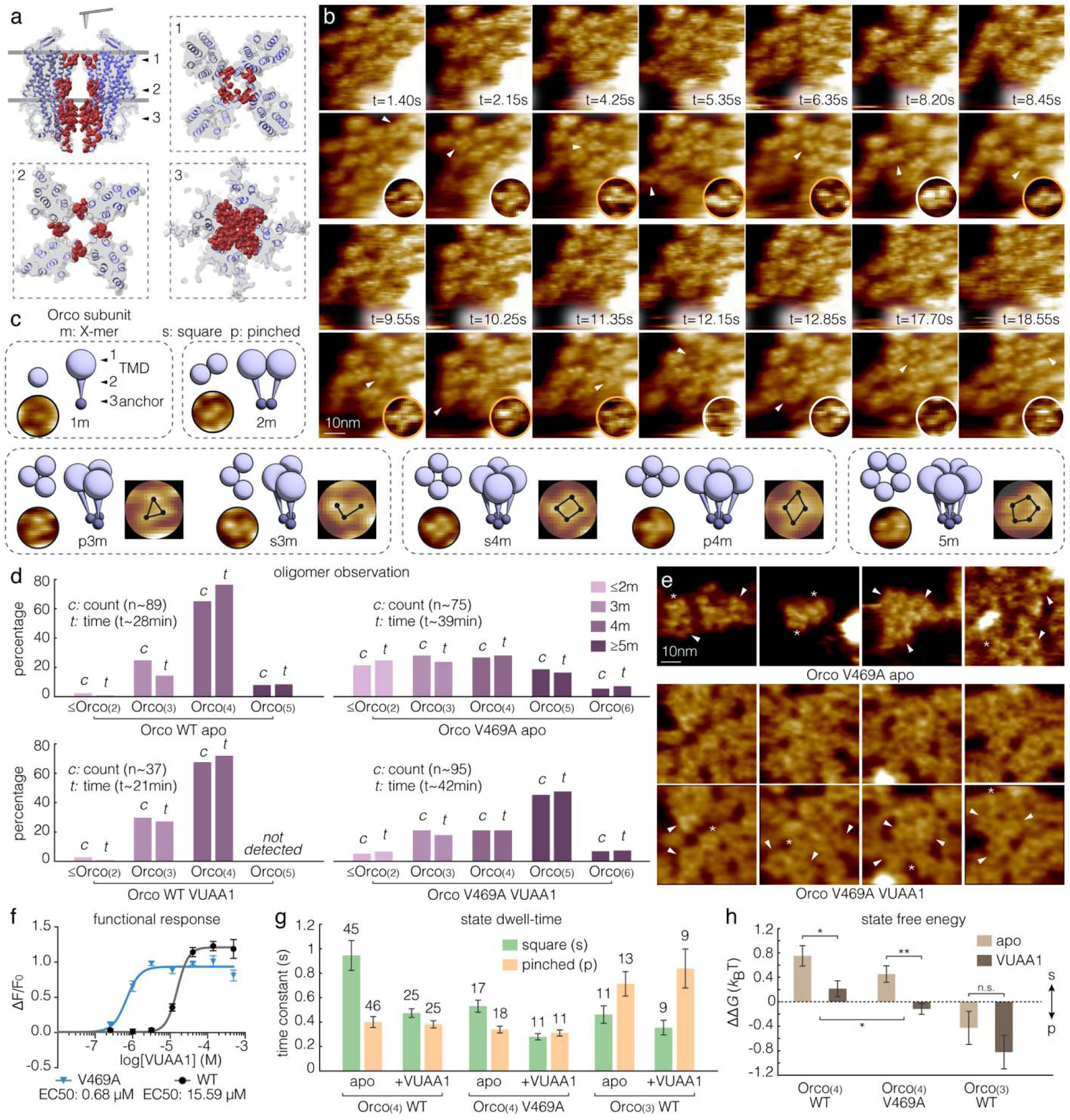
Non-canonical assemblies and symmetry-broken conformations in Orco homo-oligomers. **(a)** AlphaFold3 structural model of Orco_(4)_ (quasi-identical to the cryo-EM structure but with full-length extracellular domain, **Supplementary** Figure 1a**,b**). Top left: Side view. 1 to 3: Cross-sections of Orco_(4)_ at the extracellular gate (1), transmembrane domain (TMD) (2), and cytosolic anchor-domain (3). Interface atoms within 5 Å distance to neighbor Orco protomers are shown as red spheres. **(b)** HS-AFM of membrane-reconstituted Orco receptors (**Supplementary Movie 3**). Top: Raw movie frames. Bottom: 5-frame time-averages. Imaging parameters: 50 ms/frame, 0.67 nm/pixel. White circles: different oligomer assemblies (Orco_(5)_: t = 1.40 s and t = 12.85 s; canonical Orco_(4)_: t = 2.15 s and t = 8.20 s; Orco_(3)_: t = 4.25 s, t = 6.35 s, and t = 18.55 s; Orco_(2)_: t = 17.70 s; and Orco_(1)_: t = 12.15 s). Orange circles: non-canonical Orco_(4)_ (t = 5.35 s, t = 8.45 s, t = 9.55 s, t = 10.25 s, and t = 11.35 s) and Orco_(3)_ (t = 4.25 s and t = 6.35 s), *i.e.*, square *vs* pinched states. Arrowheads indicate the corresponding single particles. **(c)** Schematics and representative single-particle images of Orco oligomer assemblies and conformational states. From left to right: monomer (*1m*); dimer (*2m*); pinched trimer (*p3m*, triangular trimer); square trimer (*s3m*, L-shaped trimer); canonical square tetramer (*s4m*, square tetramer); pinched tetramer (*p4m*, rhombus-shaped tetramer); pentamer (*5m*). **(d)** Distribution of oligomer observations by particle-lifetime and particle-count, in apo condition or in presence of agonist VUAA1 (0.1 mM), for WT and V469A Orco. Data from a total of *n*=89 (WT apo), *n*=37 (WT VUAA1), *n*=75 (V469A apo) and *n*=95 (V469A VUAA1) particles with a total observation time of *t*∼28 (WT apo), *t∼*21 (WT VUAA1), *t∼*39 (V469A apo) and *t∼*42 (V469A VUAA1) minutes were pooled for the analysis. **(e)** HS-AFM of membrane-reconstituted gain-of-function V469A Orco receptors in apo (top: raw HS-AFM frames, **Supplementary Movie 4,5**) and VUAA1 (middle: raw HS-AFM frames; bottom: 3-frame time-averages, **Supplementary Movie 7,8**) conditions. Asterisks: canonical Orco_(4)_ assemblies. Arrowheads: Orco_(3)_, Orco_(5)_ and Orco_(6)_ assemblies. **(f)** VUAA1 dose-response curves of WT and V469A Orco homo-oligomer variants from GCaMP experiments. ΔF/F_0_: Relative fluorescence changes (mean±s.e.) for the WT (Black) V469A (Blue) Orco variants (*n*=6 biological replica). Half-maximal effective concentrations EC50: WT ∼ 15.59 μM, V469A ∼ 0.68 μM. **(g)** and **(h)** Pinching dynamics analysis of Orco_(4)_ (WT and V469A) and Orco_(3)_ (WT), in both apo and VUAA1 conditions. (g) State dwell-time constants (mean±s.e.) of the square and pinched Orco. Sample sizes are given above bars. (h) Equilibrium free energy difference (ΔΔ*G =* Δ*G*_pinched_ *–* Δ*G*_square_, mean±s.e.) between square and pinched states. Two-tailed student *t*-test (for apo *vs* VUAA1) and weighted least square analysis of variance (ANOVA, for Orco_(4)_ WT *vs* V469A): *n.s.* not significant, \**P*<0.05, \*\**P*<0.01.

To explore the functional relevance of the oligomer plasticity, we analyzed a gain-of-function Orco V469A mutant (**Figure 2e**), that has a ∼20-fold more sensitive apparent dose response to VUAA1 (**Figure 2f**, **Supplementary Figure 2**). In contrast to WT, we observed that ∼70-75% of the assemblies were non-tetrameric in the Orco V469A mutant (**Figure 2d,e**, Orco V469A apo**, Supplementary Movies 4,5, Supplementary Table 2**). Notably, Orco_(3)_ and Orco_(4)_ had comparable contribution to the overall population (∼25-30%) and thus had negligible equilibrium energy difference. Besides, lower-order (Orco_(1)_ and Orco_(2)_) and higher-order (Orco_(5)_ and Orco_(6)_) oligomers were also prevalent.

### Square and pinched state dynamics in Orco homo-oligomers

All cryo-EM structures of the OR family showed a uniform four-fold symmetric (Orco_(4)_^6^ and OR_(4)_^13^) or pseudo-four-fold symmetric (Orco_(3)_/OR_(1)_^15,16^) arrangement, with no evidence of conformational variability. Nevertheless, imaging Orco homo-oligomers with HS-AFM at 20 frames per second, we found that alongside receptors with canonical square arrangement (**Figure 2b**, white circles), many Orco_(4)_ displayed transiently a rhombus-shaped configuration, characterized by one notably elongated axis (**Figure 2b**, orange circles). Likewise, Orco_(3)_ comprised L-shaped and triangular subunit configurations (**Figure 2b**), the former resembling a tetramer missing one subunit, the latter resembling an equilateral triangle. Therefore, we categorized these oligomeric arrangements, with respect to their protomer-protomer interface, into two major states (**Figure 2c**): a ‘square’ state, including square tetramers (**Figure 2c**, *s4m*) and L-shaped trimers (**Figure 2c**, *s3m*), and a ‘pinched’ state, comprising rhombus-shaped tetramers (**Figure 2c**, *p4m*) and triangular trimers (**Figure 2c**, *p3m*). These states were computationally classified based on their inter-subunit angles (**Methods**, **Supplementary Figure 3**).

Next, we tracked single molecules for square-to-pinched and reverse transitions and quantified the ‘pinching’ dynamics. Square and pinched states had lifetimes between ∼1 s and ∼0.4 s (**Figure 2g**, Orco_(4)_ WT), and thus rate constants of ∼1 s^-1^ and ∼2.5 s^-1^, suggesting that the two states are separated by a high energy barrier. Considering an Arrhenius prefactor of ∼10^9^ s^-1^ for proteins^25^, we estimate an energy barrier in the range of ∼18-21 *k*_B_T, according to Transition State Theory.

For Orco_(4)_, the square (*s4m*) state represented the more stable configuration as compared to the pinched state (*p4m*), with an equilibrium energy difference (ΔΔ*G* = Δ*G*_pinched_ – Δ*G*_square_) of ∼0.8 *k*_B_T (**Figure 2h**, Orco_(4)_ WT). In contrast, Orco_(3)_ favored the pinched state (*p3m*), with an equilibrium energy difference of ∼ –0.4 *k*_B_T (**Figure 2h**, Orco_(3)_ WT), likely to avoid the energy penalty of exposing the hydrophilic residues in its central pore to the hydrophobic lipid tails when adopting the L-shaped state (*s3m*).

Addition of the synthetic Orco agonist VUAA1^6,26^ barely altered the assembly distribution of Orco oligomers in WT (**Figure 2d**, Orco WT VUAA1**, Supplementary Figure 4, Supplementary Movie 6, Supplementary Table 1**) but shifted the oligomer distribution to higher-order assemblies in the V469A gain-of-function mutant (**Figure 2d,e**, Orco V469A VUAA1, **Supplementary Movies 7,8**, **Supplementary Table 2**), where Orco_(5)_ became the dominant species (40-45% of all populations). VUAA1 binding also changed conformational dynamics by shortening the square state lifetime of Orco_(4)_ in both apo and VUAA1 conditions, shifting the equilibrium energy difference to ∼0.2 *k*_B_T and thus destabilizing the square state (**Figure 2g,h**). In contrast to WT, the V469A mutant was more prone to pinching, with shorter-lived square and pinched states. Taken together, the observed oligomer plasticity and pinching dynamics are modulated by ligand binding and function-altering mutation.

### Pinched conformation facilitates oligomer assembly changes

Transitions between all major oligomeric assemblies (Orco_(3)_, Orco_(4)_ and Orco_(5)_) were captured in real-time, including reversible rearrangement events (**Figure 3a-g**, **Supplementary Movies 9**-**15**). These oligomeric rearrangements occurred at timescales ranging from a few seconds to tens of seconds, corresponding to an estimated energy barrier of ∼20-25 *k*_B_T.

**Figure 3.**
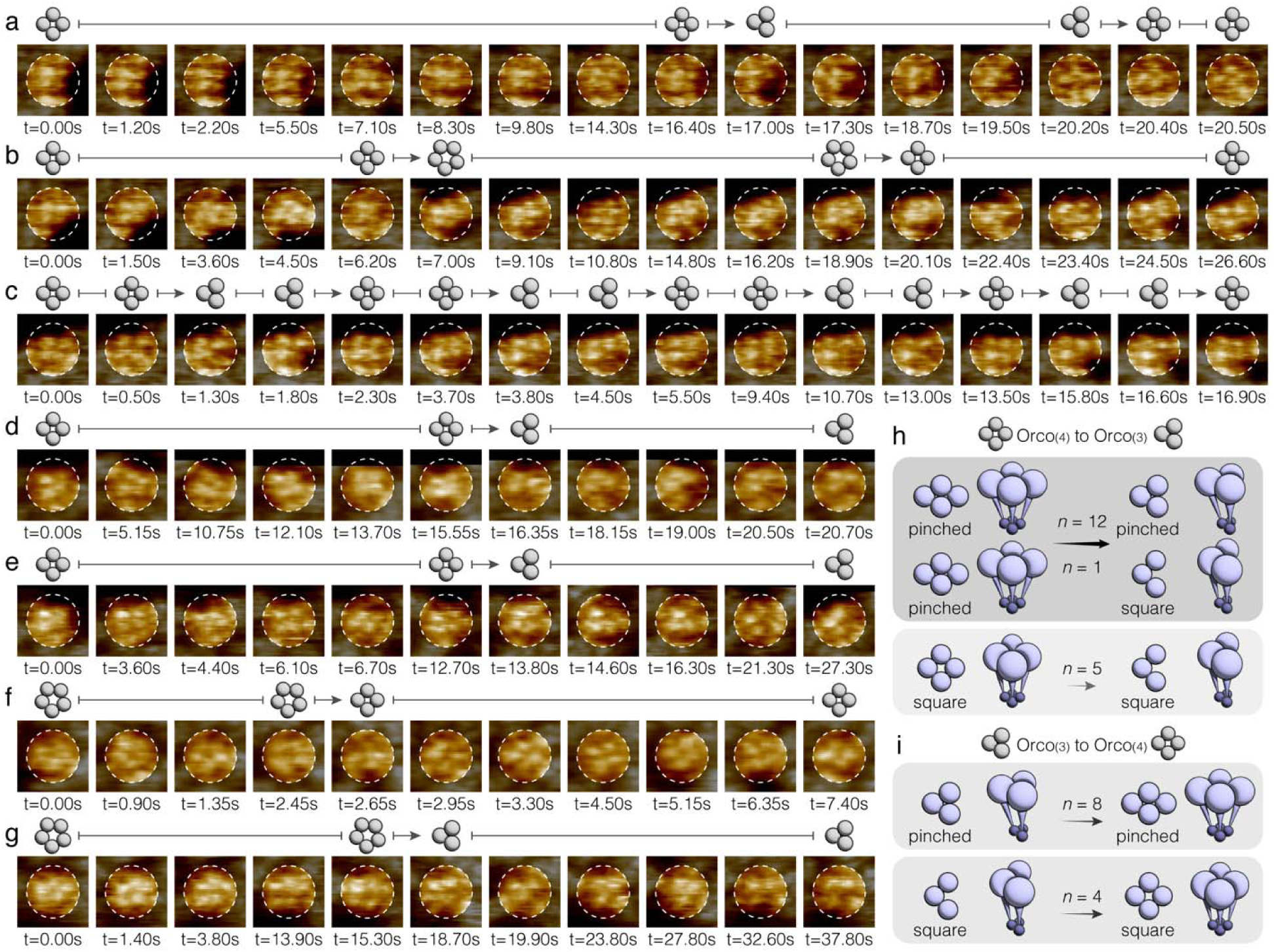
Reversible oligomeric transitions between Orco assemblies. **(a)** to **(g)** HS-AFM image series of single Orco homo-oligomers undergoing assembly rearrangements, including reversible transitions between Orco_(4)_ and Orco_(3)_ (a, **Supplementary Movie 9**) and between Orco_(4)_ and Orco_(5)_ (b, **Supplementary Movie 10**) assemblies. Frequent transitions between Orco_(4)_ and Orco_(3)_ assemblies (c, **Supplementary Movie 11**). Slow transitions from Orco_(4)_ to Orco_(3)_ (d and e, **Supplementary Movies 12,13**), and from Orco_(5)_ to Orco_(4)_ (f, **Supplementary Movie 14**) and from Orco_(5)_ to Orco_(3)_ (g, **Supplementary Movie 15**). **(h)** and **(i)** Oligomer assembly rearrangement event statistics and their associated conformational states (square vs. pinched) from Orco_(4)_ to Orco_(3)_ (h) and from Orco_(3)_ to Orco_(4)_ (i). The thickness of the arrows indicates rearrangement likelihood.

Next, we examined how oligomer assembly rearrangements are coupled to the pinching dynamics of Orco receptors (**Figure 3h,i**). Among 18 Orco_(4)_-to-Orco_(3)_ observations, 13 originated from the pinched tetramer state, yielding a conditional probability *P*_(p4m | 4m-to-3m)_ ∼ 72% (**Figure 3h**). In the case of Orco_(3)_-to-Orco_(4)_ rearrangements, 8 out of 12 events emerged from the pinched trimer state, giving *P*_(p3m | 3m-to-4m)_ ∼ 67%. To evaluate how much pinching actively influences these rearrangements, we compared these values to the baseline occurrence of pinched conformations in the overall population, *P*_(p4m)_ ∼ 32% for tetramers and *P*_(p3m)_ ∼ 60% of trimers. Applying Bayes’ theorem, we estimated that Orco_(4)_-to-Orco_(3)_ assembly rearrangements are ∼2.25× more likely to occur when the tetramer is in the pinched state (**Methods**). On the other hand, pinched trimers are more abundant than L-shaped trimers, therefore the corresponding transition gain for Orco_(3)_-to-Orco_(4)_ due to pinching is barely above expectation (∼ 1.12×). These results indicate that pinching promotes oligomer assembly rearrangements in Orco receptors.

### Oligomer plasticity in Orco/OR heteromeric receptors

In most insects, ORs function as hetero-tetramers, with a recently proposed Orco_(3)_/OR_(1)_ stoichiometry^15,16^. To investigate this functional complex, we shifted the focus to membrane-reconstituted Orco/OR heteromeric receptors. Since WT OR28 lacks an extracellular protrusion, its topography is indistinguishable from the lipid membrane and thus undetectable in HS-AFM experiments. Therefore, to make OR detectable by HS-AFM, we designed an OR28e16 subunit by introducing a short peptide of 16 amino acids into the second extracellular loop of OR28, distant from any Orco/OR inter-subunit interface. According to AlphaFold3, this peptide extends the loop by ∼0.5 nm above the membrane (**Figure 4a**, **Methods**). The Orco/OR28e16 mutant remained functional, albeit with a moderate reduction of activity (**Supplementary Figure 5**). Because of the small size of the engineered extracellular loop, the OR28e16 subunit appeared in HS-AFM as a faint protrusion, with a height ∼2-3 Å lower than the rigid extracellular beta-sheets of Orco subunits (**Figure 4b, Supplementary Movie 16,17**).

**Figure 4.**
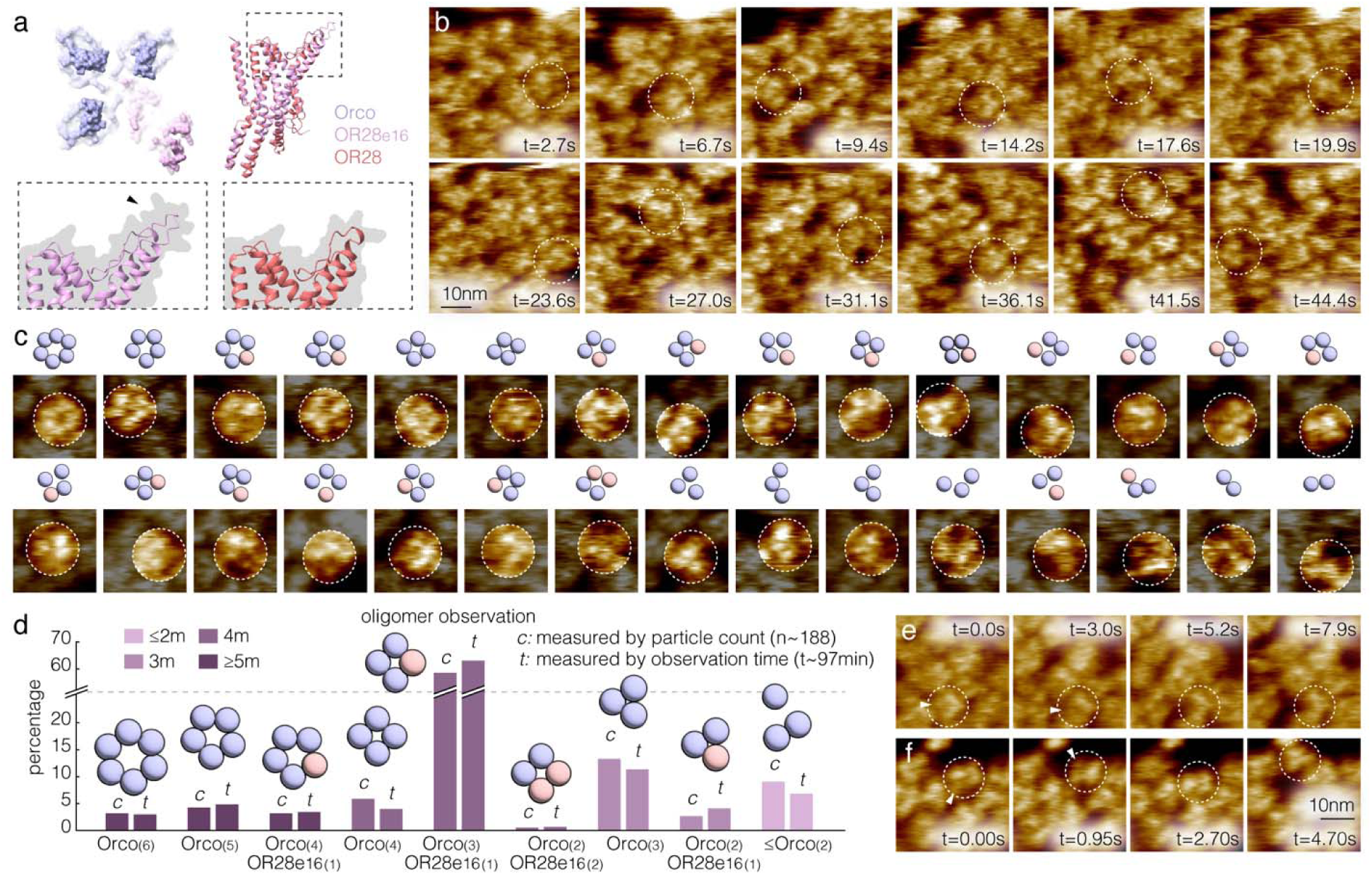
Oligomer plasticity in Orco/OR heteromeric receptors. **(a)** AlphaFold3-predicted structure of Orco_(3)/_OR28e16_(1)_ receptor. OR28e16 subunit (purple) has a short peptide insert of 16 amino acids in the second extracellular loop of OR28 (red), distant from the Orco/OR interface. Black arrowhead: Extra density in the OR28e16 subunit permits OR detection by the HS-AFM tip. **(b)** HS-AFM of membrane-reconstituted Orco/OR28e16 receptors (**Supplementary Movie 17**). Imaging parameters: 100 ms/frame, 0.5 nm/pixel. White circles: Orco_(3)_/OR28e16_(1)_ hetero-tetramers, where the OR28e16 subunit appeared as a faint but clearly detectable protrusion, with a height of ∼2-3 Å lower than the Orco subunits. **(c)** HS-AFM single-particle analysis of the Orco/OR28e16 oligomers. Top: Cartoon schematics of the oligomer configuration with Orco (blue) and Or28e16 (red) subunits. Botton: HS-AFM single-particle images. **(d)** Population distribution of the Orco/OR28e16 oligomers. Data from a total of *n*=188 different particles (*c*: particle-count) with a total observation time of ∼97 minutes (*t*: particle-lifetime) were pooled for the analysis. **(e)** and **(f)** Transitions from Orco_(3)_/OR28e16_(1)_ hetero-tetramers to Orco_(3)_ homo-trimers (**Supplementary Movies 18,19**). Dashed circle: Particle undergoing subunit rearrangement. White arrowhead: OR28e16 subunit.

The OR28e16 mutant allowed an in-depth investigation of the assembly dynamics of the Orco/OR heteromeric receptors (**Figure 4c,d**). Akin to Orco homomers, while the canonical state of Orco/OR28e16 is tetrameric, the receptors displayed significant oligomer plasticity. Indeed, ∼30-35% of the oligomers were non-tetramers (**Figure 4d**, **Supplementary Table 3**), comparable to the Orco homo-oligomer distribution (**Figure 2d**). Among the tetramer population, ∼90% were Orco_(3)_/OR_(1)_ heteromeric receptors (∼60% of all oligomers). Almost all observed Orco/OR heteromeric receptors contained only one OR subunit (>99%), regardless of their oligomeric state. These include Orco_(4)_/OR_(1)_ hetero-pentamers (∼3% of all oligomers), Orco_(3)_/OR_(1)_ hetero-tetramers, and Orco_(2)_/OR_(1)_ hetero-trimers (∼3% of all oligomers), with the sole exception of one observation of an Orco_(2)_/OR_(2)_ hetero-tetramer.

Transitions among different oligomeric assemblies were also captured in the Orco/OR28e16 sample, as we observed the dissociations of the OR subunit from Orco_(3)_/OR28e16_(1)_ resulting in a Orco_(3)_ homo-trimers (**Figure 4e,f, Supplementary Movies 18,19**). Besides, ∼30-35% of the observed molecules were Orco homo-oligomers, including Orco_(6)_ (∼3% of all oligomers), Orco_(5)_ (∼5% of all oligomers), Orco_(4)_ (∼5% of all oligomers), Orco_(3)_ (∼12% of all oligomers), and lower-oligomers (Orco_(2)_ and Orco_(1)_, ∼8% of all oligomers). Since we employed an OR-specific affinity purification strategy (**Supplementary Figure 5, Methods**), the detection of ∼30-35% OR-less oligomers further indicates the occurrence of substantial subunit rearrangement in Orco/OR receptors after membrane reconstitution and during HS-AFM imaging experiments (**Figure 4d**).

Interestingly, if (i) trimers predominantly arise from the dissociation of one subunit from the canonical Orco_(3)_/OR28e16_(1)_ hetero-tetramers, and (ii) each subunit is equally likely to dissociate, one would expect to observe 3× more Orco_(2)_/OR28e16_(1)_ hetero-trimers than Orco_(3)_ homo-trimers. In contrast, Orco_(3)_ outnumbered Orco_(2)_/OR28e16_(1)_ by ∼2.5-5 fold (**Figure 4d**, **Supplementary Table 3**). This observation indicates that OR is far less stable than Orco in the hetero-tetramers, ∼7.5-15× more likely to dissociate from the canonical Orco_(3)_/OR28e16_(1)_ hetero-tetramers, corresponding to an estimated energy difference of ∼2-3 *k*_B_T.

### Ligand binding modulates Orco/OR pinching kinetics

Ligand addition (2,4,5-trimethylthiazole, TMT, a known odorant ligand of OR28) was reported to induce minor conformational changes in Orco/OR28, as revealed by cryo-EM (**Figure 5a,b**). We therefore investigated the effect of ligand binding on the assembly and conformational dynamics of the native functional Orco/OR28 receptors. To this end, we analyzed the spatial arrangement of the three visible Orco subunits and used this information to infer the WT Orco/OR28 configurations under both apo (**Figure 5c, Supplementary Movie 20**) and ligand-bound (**Figure 5d, Supplementary Movie 21**) conditions. In HS-AFM overview imaging, we observed that >90% of the single molecules featured three membrane-protruding Orco subunits. Although the fourth position in the complex (**Figure 5e**, red) could either be an invisible OR28 subunit or an empty space, according to the Orco/OR28e16 results, the probability of a molecule with three Orco subunits to contain an OR is ∼80-85% (**Figure 4d**, percentage of Orco_(3)_/OR28e16_(1)_ among all complexes that have three Orco subunits). As in the Orco/OR28e16 analysis, we also observed non-canonical oligomers in the WT Orco/OR28 sample and captured active Orco subunit rearrangements (**Supplementary Figure 6, Supplementary Movie 22**).

**Figure 5.**
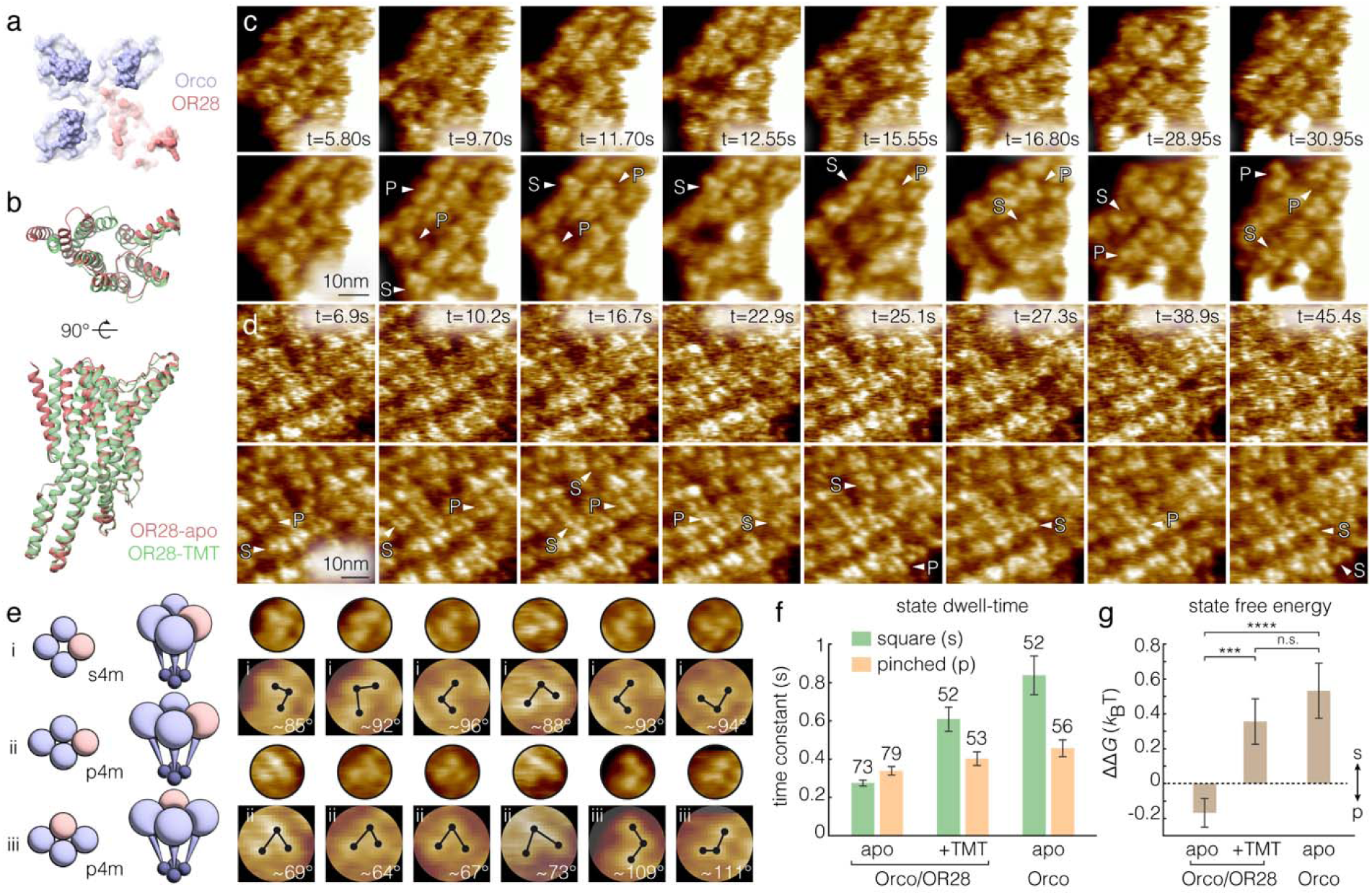
Conformational dynamics of Orco/OR28 hetero-oligomers. **(a)** Surface representation of AlphaFold3 structural model of the Orco_(3)_/OR28_(1)_ hetero-tetramer (with full-length extracellular domain). **(b)** Superimposition of OR28 subunit in apo and ligand-bound (2,4,5-trimethylthiazole, TMT) structures within the hetero-tetramer. Note the pore helix rearrangement associated with ligand-binding. **(c)** and **(d)** HS-AFM imaging of membrane-reconstituted Orco/OR28 receptors in apo (c, **Supplementary Movie 20**) and ligand-bound (d, **Supplementary Movie 21**) conditions (5mM TMT). Top: Raw movie frames. Bottom: 5-frame time-averages. Imaging parameters: 50 ms/frame, 0.67 nm/pixel (c) and 100 ms/frame, 0.5 nm/pixel (d). **(e)** Schematics and representative single-particle images of Orco/OR28 oligomers. Position in red is either empty or occupied by an OR28 subunit. Orco inter-subunit angle measurement (highlighted) is a characteristic feature that distinguishes pinched and square conformations. **(f)** and **(g)** Pinching dynamics analysis of Orco/OR28 single particles. (f) State dwell-time constants (mean±s.e.) of the square and pinched Orco/OR28. Sample sizes are given above bars. (g) Equilibrium free energy difference (ΔΔ*G =* Δ*G*_pinched_ *–* Δ*G*_square_, mean±s.e.) between the square and pinched conformations. From left to right: Orco/OR28 assemblies (with three Orco subunits clearly resolved) in apo and ligand-bound (TMT) conditions. For comparison: Orco homo-oligomers (including Orco_(4)_ and Orco_(3)_). Two-tailed student *t*-test: *n.s.* not significant, \*\*\**P*<0.001, \*\*\*\**P*<0.0001.

We measured the Orco inter-subunit angles to assess their spatial arrangement (**Figure 5e**, blue) and confirmed the presence of both square (**Figure 5c**, arrowhead *S*, **Figure 5e**, top, *s4m*) and pinched (**Figure 5c**, arrowhead *P*, **Figure 5e**, bottom, *p4m*) states in the Orco/OR28 receptors. Among the pinched state, we can distinguish two subtypes: one where the three Orco subunits form a triangle with inclusion angle <90° (∼70%), and the other with an angle >90° (∼30%). We quantified the dwell-times of the square and pinched states and estimated their equilibrium energy difference under both apo and ligand-bound conditions (**Figure 5f,g**).

Surprisingly, under apo condition, square and pinched Orco/OR28 receptors had similar lifetimes (**Figure 5f**, left) and thus free energy levels (**Figure 5g**, left), with a slight preference (∼ –0.2 *k*_B_T) for the pinched state. In contrast, in the presence of odorant (TMT), Orco/OR28 receptors exhibited a longer-lived square state with a dwell-time constant ∼600 ms, substantially longer than that observed in apo condition (**Figure 5f**, center). Thus, ligand-binding yielded a stronger preference for the square state with an equilibrium energy difference of ∼0.4 *k*_B_T (**Figure 5g**, center), comparable to Orco homo-oligomers. Since the lifetimes of the pinched state remained roughly consistent across all conditions, TMT binding induced an evident lowering of the square state energy level, by ∼0.6 *k*_B_T, thus reshaping the equilibrium energy landscape and restoring square-state stability comparable to Orco homo-oligomers (**Figure 5f,g**, right). Therefore, ligand binding modulates the pinching kinetics of Orco/OR complexes.

Beyond the single-molecule conformational dynamics, ligand-binding also altered the mobility of insect ORs within clusters (**Supplementary Figure 7**, **Methods**). In apo conditions, Orco/OR28 molecules displayed ∼2× faster lateral diffusion (*D* ∼2 nm^2^s^-1^) compared to the ligand-bound condition (*D* ∼0.8 nm^2^s^-1^) or Orco homo-oligomers (*D* ∼1.2 nm^2^s^-1^) (compare **Supplementary Movies 20** to **3,21**). In contrast, DMSO had negligible effect on the diffusive behavior (**Supplementary Figure 7d**) nor clustering, which precluded the possibility that membrane properties were altered by inorganic solvents of the receptor ligands. The observation that ORs in the presence of odorant ligand favored static cluster formation within membranes, rather than diffusing within densely packed regions, suggests the presence of protein-protein interactions of weak-to-intermediate strength (single digit *k*_B_T). Thus, the observed shift in lateral mobility of molecules, possibly related to the changes in the relative stability between square and pinched states of the receptors, may alter inter-molecular interactions and protein clustering. This observation may have relevance for the dendritic membranes of olfactory neurons, where ORs are natively enriched^10^.

### Conformational substates of a representative Orco homo-tetramer

To gain deeper kinetic insights into the pinching mechanism, we tracked the conformational dynamics of a representative Orco_(4)_ at enhanced temporal resolution (50 frames per second, 0.5 nm/pixel) for ∼80 s (**Figure 6a**, **Supplementary Movie 23**). From this single-molecule tracking, we extracted a dataset of 3,932 single-particle observations and implemented an autoencoder-based (AE)^27^ neural network for unsupervised particle classification and analysis (**Figure 6b,c**, **Supplementary Figure 8, Methods**).

**Figure 6.**
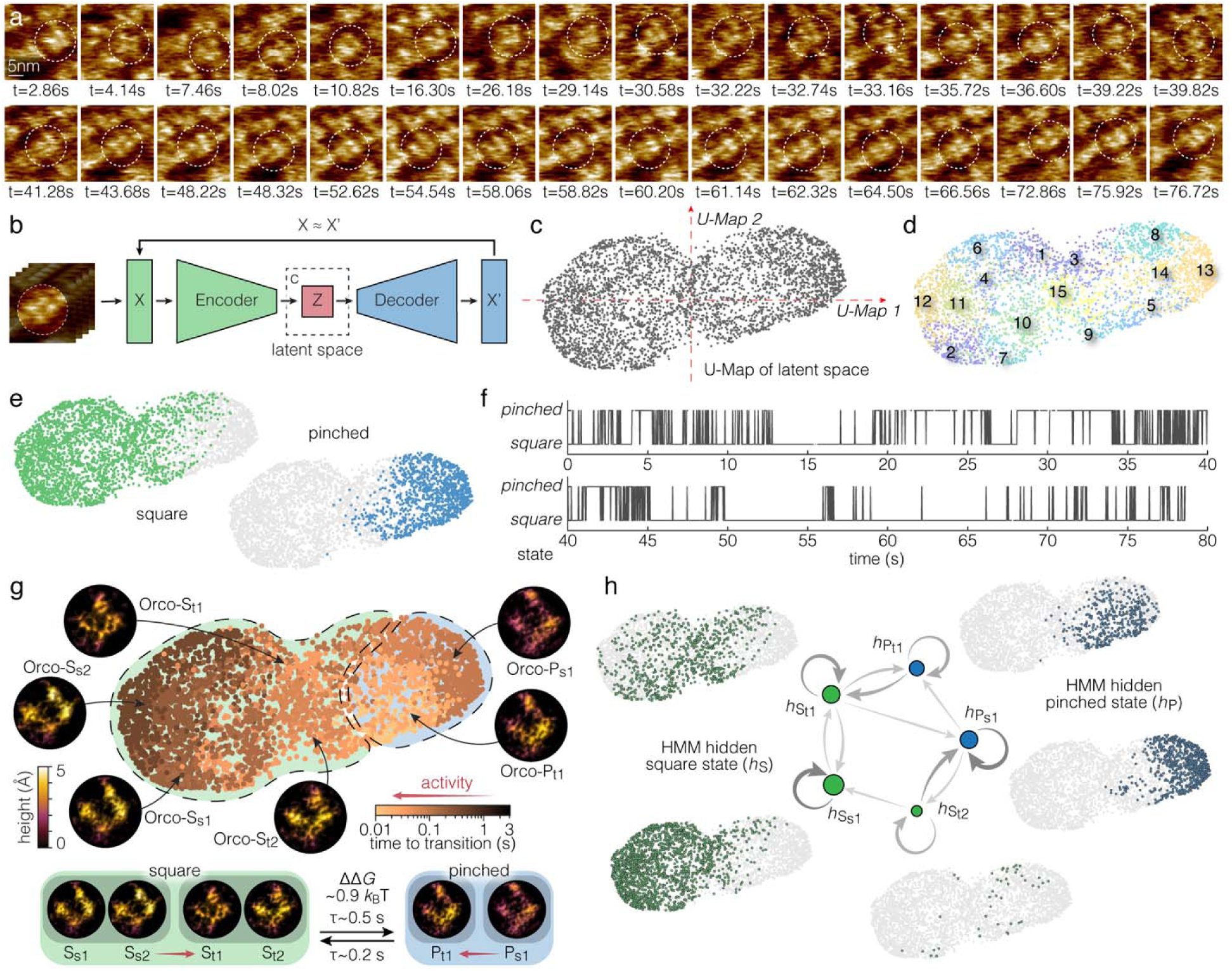
Single-molecule conformational dynamics in an Orco homo-tetramer. **(a)** Selected HS-AFM movie frames from high-resolution tracking of an Orco_(4)_ (dashed circle, **Supplementary Movie 23**). Imaging parameters: 20 ms/frame, 0.5 nm/pixel. **(b)** Implementation of an autoencoder-based (AE) neural network for HS-AFM single-particle analysis (**Supplementary** Figure 8, **Methods**). **(c)** Distribution of HS-AFM single particles in AE latent space. A U-map dimension reduction method was used to project the single-particle distribution to 2D for visualization. **(d)** Classification of HS-AFM single particles. A total of 15 conformational classes were obtained from 3,932 HS-AFM observations, based on their distribution in the AE latent space (c). **(e)** Two-state (square and pinched) conformational assignment of HS-AFM single particles, depending on the protomer arrangement in LAFM maps. **(f)** Two-state conformation-time trace of the Orco_(4)_. Multiple kinetic modes were observed from this single-molecule tracking, including time intervals during which the square state was favored (*e.g.*, t ∼ 45s to t ∼ 78s), intervals when the pinched state was favored (*e.g.*, t ∼ 20s to t ∼ 35s), and intervals characterized by rapid state interconversion (*e.g.*, t ∼ 0s to t ∼ 13s, t ∼ 35s to t ∼ 45s). **(g)** Conformation-activity analysis of the Orco_(4)_ (**Methods**), integrating the ‘time-to-transition’ measurement (**Supplementary** Figure 9) and the conformational landscape from AE neural network (c). ‘Time-to-transition’ is defined as the shortest temporal distance to a square-pinched conformational transition in the two-state conformation-time trace. Within the two major states (pinched and square), six substates were identified to contribute to the pinching transitions of this Orco_(4)_, including two stable square states (*S_s1_* and *S_s2_*, ‘time-to-transition’ >1 s, dark brown data points), two transient square states (*S_t1_* and *S_t2_*, ‘time-to-transition’ ∼0.1 s, intermediate brown data points in the left part of the conformational landscape), one stable pinched state (*P_s1_*, ‘time-to-transition’ ∼0.1-1 s, intermediate brown data points in the right part of the conformational landscape), and one transient pinched state (*P_t1_*, ‘time-to-transition <0.1 s, light brown data points in the right part of the conformational landscape). **(h)** Hidden Markov modeling (HMM) of HS-AFM single particle conformation-time trace (f). An HMM of at least five hidden substates is able to account for the two-state (square and pinched, observable states) conformation-time trace of the Orco_(4)_. This model includes three square (*h*_Ss1_, *h*_St1_ and *h*_St2_, green) and two pinched (*h*_Ps1_ and *h*_Pt1_, blue) states (**Methods**). The thickness of the arrows indicates the transition probabilities.

Based on the particle distribution in the AE latent space, observations were clustered into local conformational groups, (**Figure 6c,d**, U-map of latent space, clusters 1-15, **Methods**), and the localization AFM (LAFM)^23,28^ strategy applied to resolve super-resolution structural details for each cluster. Based on the spatial arrangement of Orco subunits in these LAFM maps, *i.e.*, their inter-subunit angles, we further classified local conformational groups (**Figure 6d**) into square or pinched states (**Figure 6e**), revealing the complex conformational landscape of the Orco_(4)_ molecule.

### Conformational substate transitions regulate Orco pinching

To bridge OR structure, *i.e.*, the conformational identity, with activity, *i.e.*, when was the conformation recorded, we integrated the conformational classification results (**Figure 6e**) with the HS-AFM timestamp of each single particle observation, resulting in a two-state, square and pinched, conformation-time trace of Orco_(4)_ (**Figure 6f**). Interestingly, this molecule displayed multiple kinetic modes, comprising time intervals during which each state was favored and intervals characterized by rapid state interconversion. Next, we plotted for each single-molecule observation the ‘time-to-transition’ metric (**Figure 6g**, false-color scale, **Supplementary Figure 9**)^22^. Interestingly, the temporal proximity to activity was strongly coupled with the substates in the conformational landscape, as revealed by the clustering of particles with similar pinching activity (**Figure 6g**). Based on these distinct structural and dynamic properties, we resolved six local conformational substates of the Orco_(4)_ (**Figure 6g**, insets). These substates include two stable square states: *Orco-S_s1_* and *Orco-S_s2_*, two transient square states: *Orco-S_t1_* and *Orco-S_t2_*, one stable pinched state: *Orco-P_s1_*, and one transient pinched state: *Orco-P_t1_* (where long to short ‘time-to-transition’ is shown from dark to light brown). Together, transitions among these stable and transient substates account for the diverse kinetic modes observed in Orco pinching dynamics.

In a parallel approach, we applied Hidden Markov Modeling (HMM) to assess the Orco conformational kinetics. The HMM analysis revealed that a minimum of five substates, two pinched and three square, were required to reproduce the observed two-state transition behavior in the Orco_(4)_ (**Figure 6f**). Remarkably, one stable square state (*h_Ss1_*) and one stable pinched state (*h_Ps1_*) were identified, both displaying frequent self-transitions and lacking a direct mutual connection. Indeed, transitions between these stable states required passage through transient states (*h_St1_*, *h_St2_*, *h_Pt1_*). Comparison of the distribution of the HMM states with the state assignment based on the ‘time-to-transition’ metric (**Figure 6g,h**, compare locations of *Orco-S_s1_*to *h_Ss1_*, *Orco-P_s1_* to *h_Ps1_*, and other transient states in the AE latent space) validates the findings from the single-particle conformation-activity analysis. These results suggest that conformational substate transitions regulate the kinetics of the Orco pinching mechanism.

### The pinched conformation alters the channel pore geometry

Although HS-AFM imaging revealed the conformational dynamics of OR complexes, it only detected the subunit dynamics based on the extracellular domain position. This led us to wonder: What is the conformational state of the TMD and pore of the pinched or square conformations? To address this question, we used 3D-LAFM densities^23^ of stable states *Orco-S_s1_* and *Orco-P_s1_* (**Figure 6g**) as physical constraints for molecular dynamics flexible fitting (MDFF) simulations to steer an Orco_(4)_ structure (**Figure 7a**) into the corresponding square and pinched conformations (**Figure 7b-e**, **Methods**). The 3D-LAFM-MDFF approach generated structural ensembles matching the HS-AFM observations.

**Figure 7.**
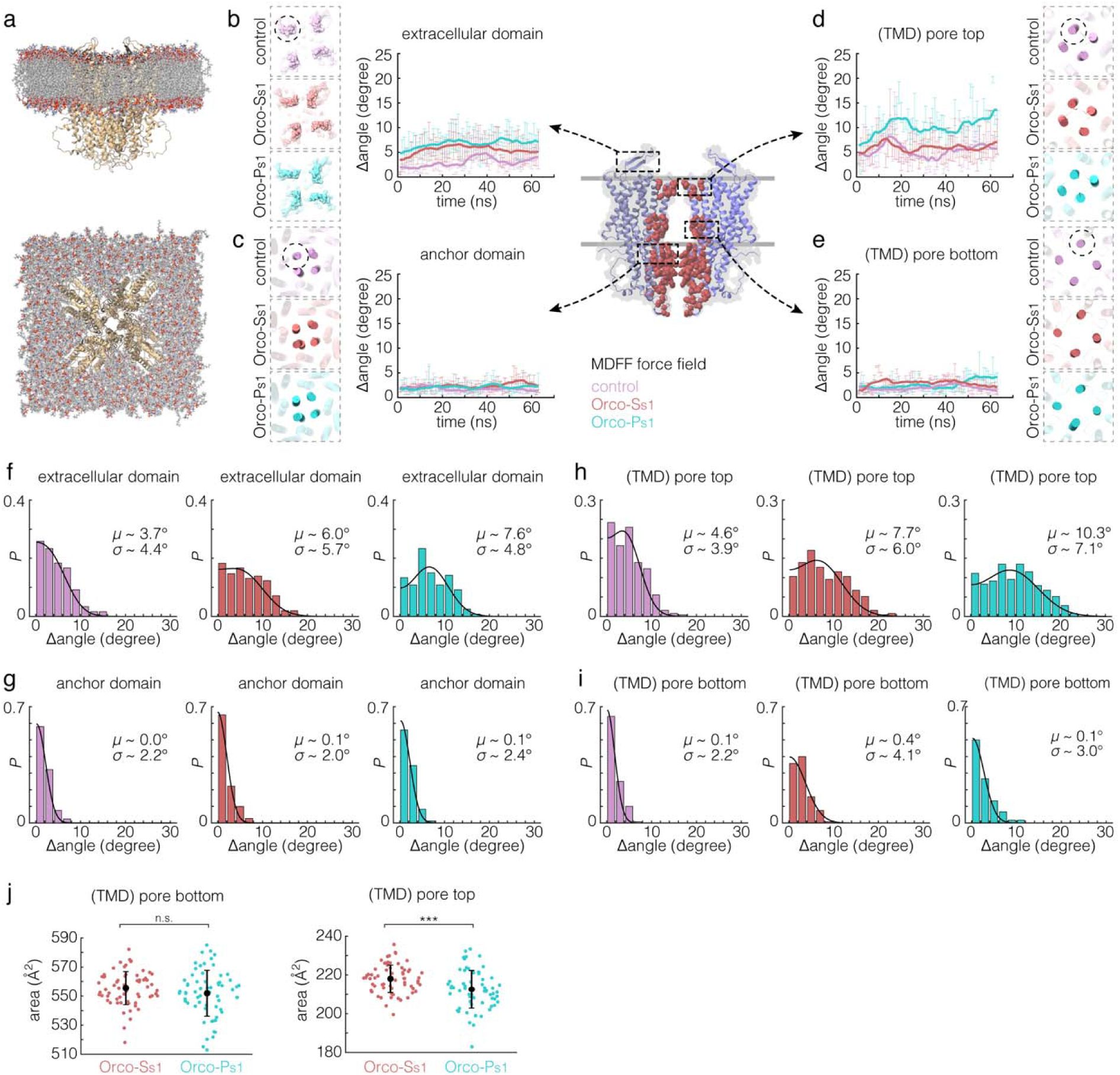
3D Localization AFM (3D-LAFM) molecular dynamics flexible fitting (MDFF) of Orco_(4)_ conformations. The 3D-LAFM maps were converted to external force fields for molecular dynamics (MD) simulations to steer an Orco_(4)_ into the corresponding HS-AFM conformations (see Figure 6), using the MDFF strategy (3D-LAFM-MDFF^23^). The stable square conformation *Orco-S_s1_* (red) and pinched conformation *Orco-P_s1_* (cyan) were used to obtain HS-AFM-derived square and pinched structural ensembles of the Orco_(4)_, along with control simulations conducted without external force fields (magenta). **(a)** Initial MD setup, box-size of 14×14 nm, with the AlphaFold3-predicted Orco_(4)_ in a lipid bilayer (**Methods)**. Three independent simulations, each 60 ns, were performed for each MDFF condition. **(b)** to **(e)** Time-evolution of collective variables (CVs) in the MD trajectories (**Methods**). These CVs encompass the extracellular domain (b), where MDFF force was applied, the anchor-domain (c), the central pore near the extracellular side (d), and the cytosolic side (e), within the transmembrane domain (TMD). The CVs characterized the inter-protomer arrangement at various locations in the protein, as measured by the deviation of the inter-protomer angles from 90° (Δangle). Measurements (mean±s.e) were taken from each structural model along the trajectories (4 Δangle per structure) at a time-step of 1 ns. Dashed boxes: Representative frames of the MD trajectories visualizing the inter-protomer arrangement. Dashed circles highlight the CVs of interest. **(f)** to **(i)** Histograms of the inter-protomer angle deviations at various locations in the protein, measured from the last 10 ns of the simulations (*N*=120 from 3 MDFF replicates for each condition). A folded Gaussian model was used to evaluate the deviation angles (**Methods**). **(j)** Central pore area measured using CVs characterizing the bottom (near cytosolic side, left) and the top (near extracellular gate, right). Two-tailed student *t*-test: *n.s.* not significant, \*\*\**P*<0.001.

We then analyzed collective variables (CVs) from the HS-AFM-derived MDFF square and pinched structural ensembles (**Figure 7f-i**). The *Orco-P_s1_* force field induced greater pore asymmetry in Orco_(4)_, with larger inter-subunit angle deviations from the initial four-fold symmetric configuration. The *Orco-S_s1_* ensemble also displayed pore asymmetry but to a much lesser extent, while the anchor-domain remained intact and symmetric across all simulation conditions. Interestingly, the enhanced pore asymmetry under the *Orco-P_s1_*force field caused a significant area reduction of ∼10 Å^2^ at the narrowest region of the pore near the extracellular gate (**Figure 7j**), which may impede ion flow. Together, the 3D-LAFM-MDFF results suggest that: (i) the observed extracellular domain pinching in HS-AFM is associated with altering central pore geometry near the extracellular gate; and (ii) the pinched conformation exhibits a more closed pore configuration.

## Discussion

Together, these findings lead us to propose a model of assembly and conformational dynamics for insect ORs in reconstituted lipid membranes (**Figure 8**): At the mesoscopic level, both Orco and OR subunits undergo membrane-diffusive protomer exchange, transitioning among various oligomeric assemblies, with the tetrameric form being the most stable. At the single-molecule level, both Orco homomeric and Orco/OR heteromeric receptors employ a pinching mechanism, switching between a symmetry-broken pinched conformation and a symmetric square conformation. Multiple substates with distinct kinetic properties further modulate the pinching behavior. In Orco/OR hetero-tetramers, ligand-binding biases the system toward the square conformation (*s4m*).

**Figure 8.**
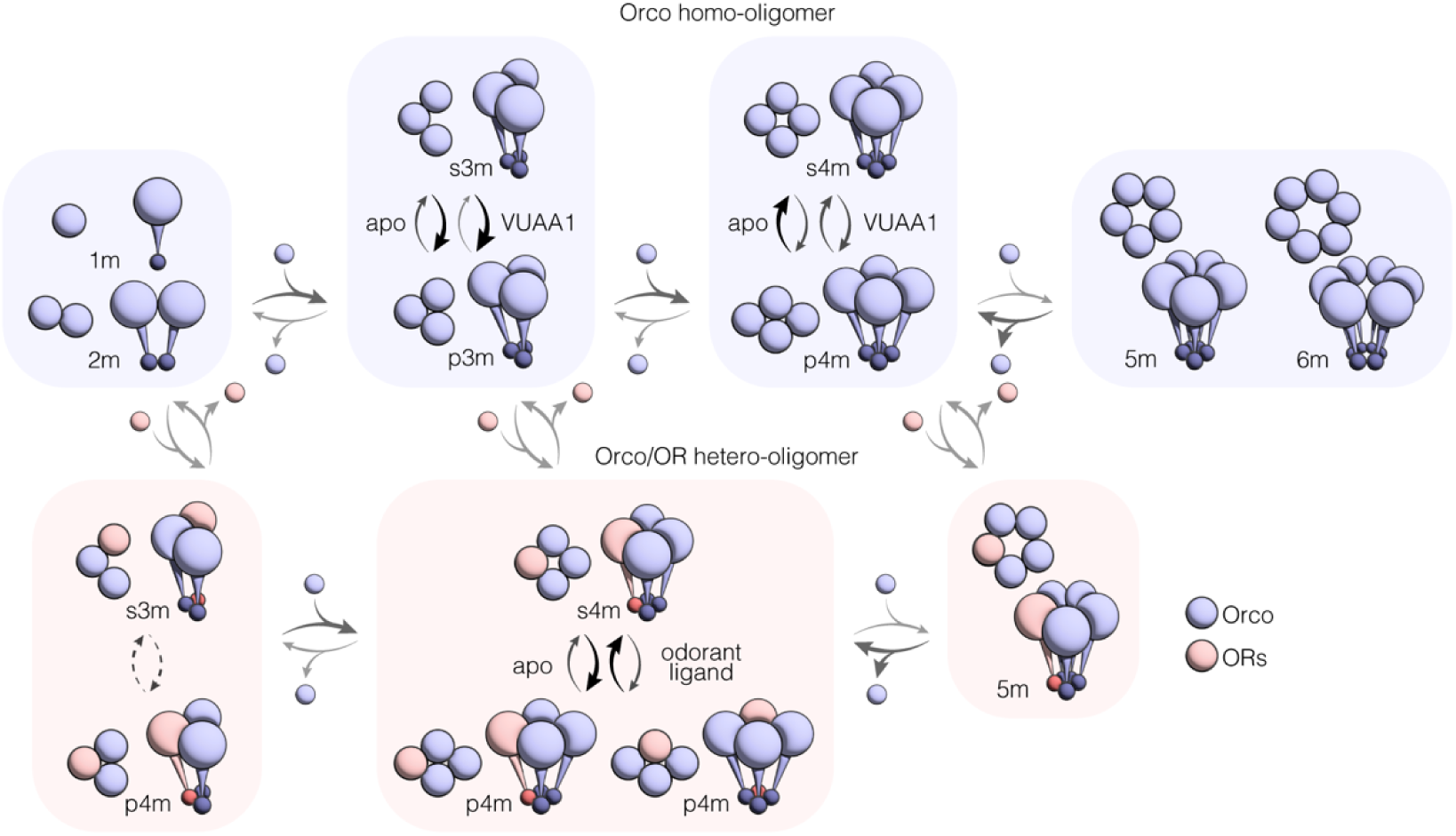
Assembly and conformational dynamics of insect ORs. Insect ORs adopt a dynamic assembly and conformational modulation mechanism. Conformational states include lower-oligomers (monomer, *1m*; dimer, *2m*), L-shaped (square) trimers (*s3m*), pinched trimers (*p3m*), square tetramers (*s4m*), pinched tetramers (*p4m*), and higher-oligomers (pentamer, *5m*; hexamer, *6m*). Among these states, Orco_(3)_/OR_(1)_ *s4m* is the canonical state. Thick arrows indicate strong transition preference.

Our findings offer valuable insights into the structure-dynamics relationship and inform subunit assembly and conformational landscape of insect OR complexes. Previous cryo-EM studies of the insect OR families have exclusively reported tetrameric assemblies^6,13,15,16^, among them the functional hetero-tetramers were shown to predominantly adopt an Orco_(3)_/OR_(1)_ stoichiometry^15,16^. Interestingly, single-molecule pull-down (SiMPull) experiment of the same purification preparation for cryo-EM showed fluorescent states consistent with fewer than 3 Orco subunits in a complex. These results were interpreted under the assumption that complexes always are tetramers and therefore suggested that other heteromeric arrangements are possible^16^. Here we show that multiple oligomeric states –with a large representation of Orco_(3)_– but also assemblies with <3 Orco without supplementing the missing subunits with OR to make up to a total of 4, also exist; which adds a layer of complexity to the interpretation of SiMPull results. In addition, MD simulations of hetero-tetramers with varying stoichiometries indicated that Orco_(3)_/OR_(1)_ represents the most thermostable assembly^16,29^.

Here, using HS-AFM single-molecule dynamic imaging, we find substantial oligomer plasticity in insect ORs, facilitated through reversible subunit exchange in membranes, and document Orco-only complexes in preparations that were OR-selectively purified (**Figure 2-4**, **Supplementary Figure 4,6**). In our experimental conditions, Orco_(3)_ was the most abundant non-canonical species, and dissociation of an OR subunit from Orco_(3)_/OR_(1)_ is highly favored over that of an Orco subunit (**Figure 4**). This suggests that second to Orco_(4)_, Orco_(3)_ represents the most stable assembly, which can rapidly incorporate a fourth subunit (Orco or OR). Since the association of an additional subunit is a second-order reaction and therefore dependent on the local density of the subunits, neurons might be able to tune the ratio of Orco-only complexes to the functional Orco/OR complexes by modulating the availability of OR subunits. These findings may also have important implications in conjunction with the recent findings that multiple ORs express in the same olfactory sensory neurons^2^, putting forward the hypothesis of OR exchange between complexes.

Besides, Orco/OR complexes were shown to display extensive conformational plasticity, transitioning between a canonical square state and a pinched state that was neither observed nor suggested in any of the cryo-EM studies (**Figure 2,5**). Using HS-AFM single-molecule structural biology analysis and MDFF, we elaborate the multi-state nature of the Orco_(4)_ pinching kinetics and find that the pinched Orco_(4)_ displays increased asymmetry and area reduction of the pore (**Figure 6,7**). Interestingly, binding of VUAA1 (Orco agonist) to Orco_(4)_ stabilized the pinched state, while binding of TMT (OR28 ligand) to Orco_(3)_/OR28_(1)_ stabilized the square state. This apparent discrepancy might be explained by the fact that VUAA1 induces conformational changes in all four Orco subunits, rendering the complex prone to pinching, whereas TMT binding perturbs only the OR28 subunit and leaves the three Orco subunits more stable within their preferred square arrangement. Previous MD simulations of Orco_(3)_/OR_(1)_ also suggested that the hetero-tetramer can adopt a non-square state, with a symmetry-broken extracellular domain and pore but a symmetric anchor-domain, comparable to the pinched state reported here and likely associated with ligand binding to the OR subunit (**Supplementary Figure 10**)^29^.

More importantly, the gain-of-function Orco mutant V469A displayed enhanced oligomer and conformational plasticity than WT, manifesting direct functional relevance of the observed dynamics. Both oligomer plasticity and pinching mechanism appear intuitively related to the Orco/OR structure, where most inter-subunit contacts are mediated by the cytosolic anchor-domain, while only few contacts are established between the subunits’ TMDs. These observations suggest that opening of the channel could be tied to assembly and conformational dynamics of not just the OR subunit, but also the Orco subunits. The consistent structural and functional features shared by insect ORs from species that diverged millions of years ago further suggest that the dynamics investigated in this study, in which the widely conserved Orco plays a vital role, are likely to be broadly relevant across the entire OR family.

Therefore, the assembly and conformational dynamics reported here represent an emerging framework of dynamic regulation in insect ORs; and may be more broadly applicable to other membrane protein systems where hetero-oligomeric complexes are the functional units. It is worth noting that HS-AFM is particularly effective for studying these processes, owing to its unique capacity to capture protein structural dynamics in real time at the single-molecule level.

## Methods

### Expression and Purification of insect odorant receptors

The Orco gene was cloned into a pEG BacMam vector with an N-terminal mCherry tag followed by an HRV 3C protease site. The wild-type OR28 gene was cloned into the same vector with an N-terminal superfolder GFP and a HRV 3C protease site. The engineered OR28 variant (OR28e16) contains a 16-residue insertion (SGGSDYKDDDDKSGGS) between Glu180 and Ser181 in the C-terminal region of extracellular loop 2. OR28e16 was identified through screening of multiple extracellular-loop epitope tag insertions using a GCaMP-based functional assay. For co-expression of OR28 and Orco, the pEG BacMam vectors were modified by inserting a tet operator (tetO) sequence downstream of the CMV promoter. For transient transfection, OR28 and Orco plasmids were mixed at a ratio of 1:2 (w/w). A total of 450 mL Expi293F inducible cells at 2.5 × 10 cells/mL were co-transfected with 0.34 mg of plasmid DNA using 0.4 mL FectoPRO and cultured at 37°C with 8% CO, shaking at 125 rpm. Protein expression was induced 48 hours post-transfection by addition of 10 mM sodium butyrate, 0.36% glucose, and 4 µg/mL doxycycline at 30°C. Cells were harvested 48 hours after induction and stored at –80°C until use. Purification of the wild-type and engineered OR28/Orco complexes followed the previously described protocol^15^.

### Lipid preparation

Lipids (1,2-dioleoyl-sn-glycero-3-phosphocholine (DOPC), 1,2-dioleoyl-sn-glycero-3-phospho-L-serine (DOPS), and Cholesterol purchased from Avanti polar lipids were solubilized in chloroform. The lipid mixture (DOPC:DOPS:Cholestrol at 8:1:1, wt:wt:wt) was dried by a gentle steam of nitrogen gas and further dried in a vacuum chamber overnight. The dried lipids were rehydrated to a final lipid concentration of 10 mg/ml, and subsequently tip-sonicated for 2 minutes to obtain small-unilamellar vesicles (SUVs).

### Reconstitution of insect odorant receptors

Purified insect odorant receptors were diluted to a final protein concentration of 0.5 mg/ml with the reconstitution buffer containing: 20 mM HEPES at pH 7.5, 150 mM NaCl, 0.01% LMNG, and 0.002% CHS. The lipid mixture (SUVs of DOPC:DOPS:Cholestrol at 8:1:1, wt:wt:wt) was supplemented to the protein reconstitution mixture at lipid-to-protein ratio (LPR) of 1.5. This mixture was allowed to equilibrate for >1 hr, after which ∼1 mg of wet Bio-Beads (Bio-Rad) per 10 μL sample was added for detergent removal. The reconstitution process could last over a few days until proteo-liposomes were harvested, with the Bio-Beads being replaced every 7-12 hours. The proteo-liposomes were checked by negative-stain electron microscopy for the presence of protein-packed vesicles of intermediate sizes (100-500 nm). Other lipid mixtures were also used for reconstitutions, including DOPC:DOPS:DOPE at 8:1:1 (wt:wt:wt), 1-palmitoyl-2-oleoyl-glycero-3-phosphocholine (POPC):DOPS:Cholesterol at 8:1:1 (wt:wt:wt), and, to analyze the potential effect of different hydrophobic mismatch, 1,2-dimyristoleoyl-sn-glycero-3-phosphocholine (14:1PC)/ 1,2-dimyristoyl-sn-glycero-3-phospho-L-serine (14:0 PS)/Cholesterol at 8:1:1 (wt:wt:wt). Similar protein behaviors were observed in all of these reconstitution conditions.

### High-speed atomic force microscopy (HS-AFM)

HS-AFM measurements were performed with an HS-AFM (RIBM) operated in amplitude modulation mode. Igor Pro version 7 was used for HS-AFM data collection. In brief, we used short cantilevers (USC-F1.2-k0.15, NanoWorld) with a nominal spring constant of 0.15 N m^−1^, a resonance frequency of ∼0.6 MHz, and a quality factor of ∼1.5 in the aqueous environment. All data were acquired at standard laboratory temperature (298 K). HS-AFM movies were aligned, flattened, and calibrated using customized ImageJ plugins (ImageJ, NIH). Both trace and retrace movies were exported and subsequently integrated to a single movie using a customized Unet-based neural network.

### HS-AFM sample preparation and physisorption

Reconstituted samples were diluted with imaging buffer (20 mM Tris-HCl at pH 7.6 and 150 mM NaCl, apo condition), of which 2 μL was deposited onto freshly cleaved mica and incubated for 8 minutes for physisorption. The excess proteo-liposomes, not physiosorbed to mica, were rinsed with the imaging buffer. Either trimethylthiazole (TMT, 5 mM, Sigma) or - for control experiments - dimethyl sulfoxide (DMSO,

10 mM, Sigma) was added to the AFM fluid chamber during imaging for the ligand-bound or DMSO conditions, respectively. Given the observed lateral mobility of the complexes in the membrane, physisorption minimized the physical interaction with the mica support.

### HS-AFM single-molecule tracking

HS-AFM single-molecule analyses were performed using customized MATLAB (MathWorks) scripts. For single-molecule tracking, HS-AFM movies were first time-averaged, from which particles were picked using customized ImageJ plugins. Coordinates of picked particles, along with their corresponding timestamps and particle identifiers, were exported and used as input for MATLAB. Single-particle coordinates and timestamps were reorganized to generate raw single-particle traces, which were then manually corrected to ensure accurate tracking of fast-diffusing molecules. These raw single-particle traces were subsequently used as templates to reconstruct the fine single-particle traces from the original non-averaged HS-AFM movies. This operation was essential for accurate single-particle analysis and dwell-time estimation, free of bias introduced by time-averaging.

### Oligomeric assembly analysis

For single-particle oligomeric assembly analysis, Orco subunits were located based on their membrane protrusion features in individual particle image. This allowed the classification of particles into distinct oligomeric states. In the case of Orco homomeric and Orco/OR28e16 heteromeric receptors, the population distribution of different oligomeric assemblies was quantified, and the equilibrium energy differences were estimated by taking the natural logarithm of the population ratios. Assemblies with less than three Orco subunits (dimers and monomers) were rarely detected due to their weak topological features, and their real populations may be underestimated in our analysis. Note that HS-AFM imaging was performed ∼5-10 minutes after the 8-minute physisorption, and for another ∼10 minutes for ligand-bound conditions after ligand addition. Thus, the observed oligomer populations (total wait time after sample addition: 15-20 minutes for apo, and 25-30 minutes for ligand-bound conditions) should represent the equilibrium distributions.

### Square and pinched state analysis

For single-particle pinching analysis (**Supplementary Figure 3**), we focused on Orco_(3)_ and Orco_(4)_ homo-oligomers, as well as Orco/OR28 particles with three Orco subunits (either Orco_(3)_ homo-trimers or Orco_(3)_/OR_(1)_ hetero-tetramers with one invisible OR subunit). Inter-subunit angles were calculated for individual particle images and used as the criterion for state determination. For any Orco trimer set (Orco_(4)_ have four such trimer subsets), we have three cases: (1) if the largest inter-subunit angle (∼90°) is approximately equal to the sum of the two smaller angles (∼90°), the configuration was considered ‘square’. (2) If the largest inter-subunit angle (∼60°) is approximately equal to 0.5× the sum of the two smaller angles (∼60°), *i.e.*, three inter-subunit angles are approximately identical (∼60°) (case 2), or (3) if the largest inter-subunit angle (∼120°) is approximately equal to 2× the sum of the two smaller angles (∼60°) (case 3), the configuration was considered ‘pinched’. For a triangle set with angles α*_1_*, α*_2_*, and α*_3_*, where α*_1_* is the largest, a sum squared error value (ε) was calculated for each of the three cases, as:

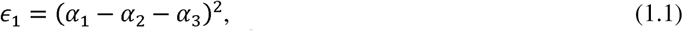

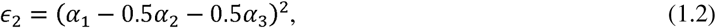

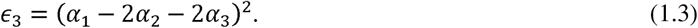

If and only if ε*_1_*is the smallest of ε*_1_*, ε*_2_*, and ε*_3_*, the particle is assigned to the square configuration. The square and pinched state dwell-time distributions were fitted with a single exponential to assess the time constants, and the equilibrium energy differences were estimated by taking the natural logarithm of the time constant ratios. Note that AFM tip may disturb proteins while imaging if imaging force is not cautiously controlled, which might lead to symmetry-broken configurations. In this case, since AFM x-axis and y-axis represent the fast and slows scanning directions, respectively, tip disturbance and thus force applied to protein should have anisotropic bias. However, there was no directional bias observed in protein pinching (**Figure 2b,c**, **Figure 5c-e**, **and Figure 6a**). Thus, it is highly unlikely that the observed pinching is a result of tip disturbance.

### Conditional probability analysis of oligomer plasticity given the pinched conformation

The conditional probability of Orco_(4)_-to-Orco_(3)_ transitions originating from the pinched tetramer state is *P*_(p4m | 4m-to-3m)_ ∼ 72%. The conditional probability of Orco_(3)_-to-Orco_(4)_ transitions originating from the pinched trimer state is *P*_(p3m | 3m-to-4m)_ ∼ 67% (**Figure 3h**). To assess to which extent pinching actively influences these rearrangements, we compared these values to the baseline occurrence of pinched conformations in the overall population, *P*_(p4m)_ ∼ 32% for tetramers and *P*_(p3m)_ ∼ 60% of trimers. In the case of the Orco_(4)_-to-Orco_(3)_ transition, the unconditional probability of a molecule undergoing such a rearrangement during the observation window is *P*_(4m-to-3m)_, and the conditional probability of this transition coming out of a pinched state is *P*_(4m-to-3m | p4m)_. Applying Bayes’ theorem, the gain in rearrangement likelihood due to pinching is *P*_(4m-to-3m | p4m)_/*P*_(4m-to-3m)_ = *P*_(p4m | 4m-to-3m)_/*P*_(p4m)_ ∼ 2.25. Similarly, we calculated *P*_(3m-to-4m | p3m)_/*P*_(3m-to-4m)_ ∼ 1.12 for the Orco_(3)_-to-Orco_(4)_ transitions.

### In-cluster mobility analysis

To estimate the mobility of single molecules within the cluster, we first calculated their stepwise displacement from the coordinates at frame *n* (*r*_n_) and frame *n*+1 (*r*_n+1_). For particle *i*, the stepwise displacement is Δ*r_n,i_*=*r*_n+1,i_-*r_n,i_*. From each frame, we calculated the mean particle displacement <Δ*r*_n_> to characterize collective motion of particles or mechanical drift during HS-AFM imaging, and then determined the drift-corrected displacement as Δ*r_n,i_*-< Δ*r_n_*> Therefore, for each HS-AFM movie with an imaging rate of Δ*t* s/frame, the mobility *D* was calculated as:

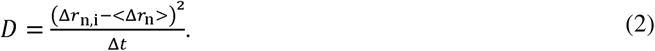

### HS-AFM single-molecule structural biology analysis

HS-AFM single-molecule structural biology analysis was performed with customized MATLAB scripts, modified from previously published protocol^22^. A dataset of 3,932 single-particle images (32×32 pixels) was constructed from a high-resolution HS-AFM tracking movie of an Orco_(4)_ (20 ms/frame, 0.5 nm/pixel, total duration 78.64 s). The single-particle images were fine-aligned using a customized ImageJ plugin. For the autoencoder (AE) neural network implementation and training, we used the MATLAB ‘trainAutoencoer’ package. AE networks are trained to project high-dimensional data onto a low-dimensional ‘latent space’ via an ‘encoder’ and subsequently reconstruct high-dimensional data through a ‘decoder’ with minimal loss (**Figure 6b**). Trained AEs are powerful in learning essential features from complex datasets in the form of latent variables and are therefore widely applied in single-particle analysis across structural biology methods^27^. By augmenting the input dataset with random rotations and translations, we trained an AE that effectively captured structural features of HS-AFM single-particle images (**Figure 6c**, U-map of latent space), while ignoring irrelevant information (**Supplementary Figure 8**, *e.g.*, scan lines, anisotropic particle orientations, and alignment errors).

A tandem framework of two AE networks was employed. First, principal component analysis (PCA) was used to roughly estimate the dimensionality of the AE latent space required to encode the input dataset (32×32=1024 dimensions). Based on this, an AE network was trained (AE1) to encode input dataset into a 55-dimensional latent space. A subsequent PCA of the AE1 latent space suggested that a 22-dimensional latent space would sufficiently capture the key characteristics of the input image. Thus, a second AE network (AE2) with a 22-dimensional latent space was trained, and its latent space was used for downstream single-molecule classification and analysis. For 2D visualization, the 22-dimensional AE2 latent space was reduced using the Uniform Manifold Approximation and Projection (Umap) method. For local particle clustering, K-means clustering was applied to the cosine similarity matrix derived from the AE2 latent space. A cluster size of ∼250 particles was used to satisfy the requirement for 3D localization AFM map construction, yielding 15 clusters from the total of 3,932 particles (**Methods**: ‘3D Localization AFM (3D-LAFM)’).

From the 3D-LAFM maps of individual clusters, we measured the spatial arrangement of the Orco subunits, based on which 15 clusters were further classified into square and pinched conformational classes. Integrating the class IDs with the HS-AFM timestamps of single particles, we constructed a two-state conformation-time trace. Notably, clusters 8 and 15 resulted in low-quality LAFM reconstructions, presumably due to limited HS-AFM image resolution. As a result, they were assigned to the nearest state in the time trace, unless the maximum frame gap exceeded 3 frames, in which case an undetermined state (*NaN*) was assigned. Subsequently, we measured the ‘time-to-transition’ metric for individual particles, which is defined as the shortest temporal distance to a square-pinched conformational transition in the two-state conformation-time trace, as an estimate of their pinching activity level.

In summary, the 22-dimensional AE2 latent space characterizes the structural properties of the particles, while the ‘time-to-transition’ metric reflects their activity properties. By integrating both structural and activity matrices, we resolved six distinct conformational substates underlying Orco pinching dynamics. These substates include two stable square states (‘time-to-transition’ >1 s): *Orco-S_s1_* and *Orco-S_s2_*, two transient square states (‘time-to-transition’ ∼0.1 s): *Orco-S_t1_* and *Orco-S_t2_*, one stable pinched state (‘time-to-transition’ ∼ 0.1-1 s): *Orco-P_s1_*, and one transient pinched state (‘time-to-transition’ <0.1 s): *Orco-P_t1_*.

Note that this analysis requires advanced particle classification, and thus it is challenging to combine datasets at the moment. Therefore, fast imaging of the same object at high spatio-temporal resolution was performed to collect a sufficient dataset. The underlying assumption is that during the imaging period (∼80 second), the molecule has extensively explored its conformational landscape, given that the pinching kinetics has sub-second state dwell-times. Though the analysis is limiting to one representative molecule, the conformational landscape provides insights into the conformational and kinetic heterogeneity of the pinching mechanism.

### 3D Localization AFM (3D-LAFM)

3D-LAFM was performed using customized MATLAB scripts, modified from previously published protocol^22^. For each single-particle cluster, images were rotationally and translationally aligned. Local maxima were identified in each image and subsequently localized to sub-pixel precision using a local image expansion strategy with a scaling factor of 50. The localized pixels and the corresponding single-particle alignment parameters were then integrated to yield the LAFM detections, which were subsequently allocated to a 3D detection stack *D*_ijk_ with a voxel size of 0.5 Å, where *i*, *j*, and *k* correspond to the *x*, *y*, and height (*z*) dimensions in the AFM images. A ‘half-bit wavelength’ (λ_hb_) was calculated as an assessment of data quality for each 3D-LAFM detection stack, derived from the Fourier shell correlation (FSC) curve between two single-particle half stacks. 3D-LAFM detection stacks had a λ_hb_ of ∼1.5-3 Å in this study. A 3D Gaussian density function was applied to transform each detection stack *D*_ijk_ to the final 3D-LAFM density map *P*_ijk_, using the computationally derived σ value equivalent to λ_hb_. All detections were included in the 3D-LAFM pipeline with no molecular symmetry applied in this study.

### Hidden Markov modeling (HMM)

HMM was performed using customized MATLAB scripts. In our study, two visible states, square and pinched, were defined, and their transitions were characterized by the two-state conformation-time trace from the HS-AFM single-molecule structural biology analysis (see main text, **Methods**: ‘HS-AFM single-molecule structural biology analysis’), from which a total of 6 conformational substates from 15 clusters were identified. Accordingly, we initially proposed an HMM with 15 hidden states as an initial guess (HMM1: 15 hidden states, 2 visible states). The maximum likelihood estimate of the transition matrix (note, the emission matrix was omitted to simplify the model) was calculated from the two-state trace using the Viterbi algorithm for this initial model. To obtain the final model, we precluded hidden states that only exhibit self-transitions (accommodating 47 particles from 6 hidden states) or no self-transitions (accommodating 31 particles from 4 hidden states), because these states may either be disconnected from the rest of the model or act solely as redundant paths.

Next, we proposed an HMM with 5 hidden states as the final model (HMM2: 5 hidden states, 2 visible states). The maximum likelihood estimate of the transition matrix, using the Baum-Welch algorithm, yielded two stable states, one square (*h*_Ss1_) and one pinched (*h*_Ps1_), that display frequent self-transitions and lacking a direct mutual connection. Transitions between these stable states required passage through transient states, including two square (*h*_St1_ and *h*_St2_) and one pinched (*h*_Pt1_). Particles attributed to the hidden states were also determined using the maximum likelihood estimation. HMM was used as an independent approach alongside the ‘time-to-transition’ metric. Both analyses reflect the activity properties of single particles and were compared to the structural properties represented in the AE latent space. The corroborative and cross-validated results from both methods highlight the multi-state nature underlying the conformational dynamics.

### 3D-LAFM molecular dynamics flexible fitting (MDFF)

3D-LAFM-MDFF was performed using customized MATLAB scripts, modified from previously published protocol^23^. MDFF force fields corresponding to stable states *Orco-S_s1_* and *Orco-P_s1_* were generated to steer an AlphaFold3-predicted Orco_(4)_ structural model into the square and pinched conformations observed in HS-AFM experiments (AFM-derived structural ensembles), respectively. In brief, 3D-LAFM density maps of these conformations were converted into MDFF force fields by aligning them to the structural model in ChimeraX (UCSF) and filling the surrounding space with a minimal background value. The resulting force fields, *U*_AFM_*-Orco-S_s1_* and *U*_AFM_*-Orco-P_s1_*, were then applied to residues 157-161 and 161-171, corresponding to the extracellular beta-sheet elements.

MDFF was performed following a standard protocol^30^. In brief, an initial simulation setup with a box size of 14 nm, where the AlphaFold3-predicted Orco_(4)_ model was embedded into a lipid bilayer matching the experimental conditions (DOPC:DOPS:cholesterol at 8:1:1, wt:wt:wt), was generated using CHARMM-GUI^31^. The system was then allowed to equilibrate following the standard six-step protocol. MDFF simulations were set up in VMD (version 1.9.4^32^) and performed using NAMD (version 2.13^33^) with the CHARMM27 force field^34^. The atomic structure was first rigid-body docked into the target *U*_AFM_, in which the extracellular beta sheets were largely submerged in the 3D-LAFM maps. To avoid overfitting and prevent structural artifacts, restraints on dihedral angles and hydrogen bonds, as well as restraints to preserve the cis/trans configuration and chirality, were applied to enforce secondary structure, following standard MDFF practice.

All simulations in this study were performed for 60 ns, comprising three cycles. Each cycle commenced with an initial energy minimization step of 10 ps, followed by a running process of 20 ns, and concluded with an additional minimization step of 10 ps. Domain restraints were applied during the first two cycles to assist large-scale structural movements (*e.g.*, the repositioning of the protein/protomer to match the 3D-LAFM map, guided by the extracellular beta-sheets). These restraints were applied to residues 59-63, 67-71, 147-151, 167-171, 188-192, 367-371, and 378-382. Three replicates were performed in each simulation condition. In control simulations, no *U*_AFM_ force field was applied. All simulations were conducted at a constant temperature of 300 K and pressure of 1.01 bar, using Langevin dynamics with a damping coefficient of 5 ps^−1^, a time step of 1 fs, and periodic boundary conditions.

### 3D-LAFM-MDFF simulation trajectory analysis

3D-LAFM-MDFF simulation trajectory analysis was performed using customized MATLAB scripts. Collective variables (CVs) were defined to characterize various structural elements, including: (1) an anchor-domain loop (residues 434-438, which harbor the majority of protomer-protomer interactions and is thus related to the stability of the oligomer), (2) the central pore loop in the TMD near the anchor-domain (residues 453-457, which define the pore integrity and is related to the ion permeation pathway), (3) the central pore loop in the TMD near the extracellular domain (residues 469-473, which define the pore integrity and is related to the ion permeation pathway), and (4) the extracellular beta sheet (residues 157-161 and 161-171, which account for tip-sample interactions and is thus directly related to the AFM conformations; and subjected to AFM-data derived force field). CV measurements from the last 10 ns of the simulations were pooled for statistical analysis of the final AFM-derived structural ensembles. Inter-protomer angles (α) were calculated based on the center-of-mass positions of the CVs, derived from the dot products of CV vector pairs connecting adjacent subunits in the tetramer (four inter-protomer angles per structure). The distribution of their deviations from 90° (Δangle = |α − 90°|) was fitted with a folded Gaussian, using the following formula:

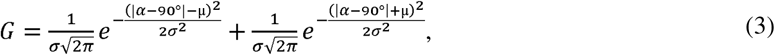

The pore area at its narrowest regions was calculated from the cross product of CV vector pairs connecting subunits along the diagonals of the tetramer.

## Author contributions

Y.J., J.M., and S.S. designed the study; J.Z. and J.M engineered the AgOR28, characterized the function, and purified all the proteins; Y.J. performed the HS-AFM experiments; Y.J. and S.S. performed the HS-AFM analyses; Y.J. designed and performed the HS-AFM single-molecule structural biology analysis; Y.J designed and analyzed the MDFF; Y.J. and Z.W. performed the MD simulations; Y.J. and S.S. wrote the manuscript; All authors edited the manuscript; S.S. supervised the study.

## Data and materials availability

All data needed to evaluate the conclusions in the paper are present in the paper and/or the Supplementary Materials. Additional data related to this paper may be requested from the authors.

