## Supplementary Information for "Assembly and Conformational Dynamics of Insect Odorant Receptors"

### Supplementary Figures

16 **Supplementary Figure 1**

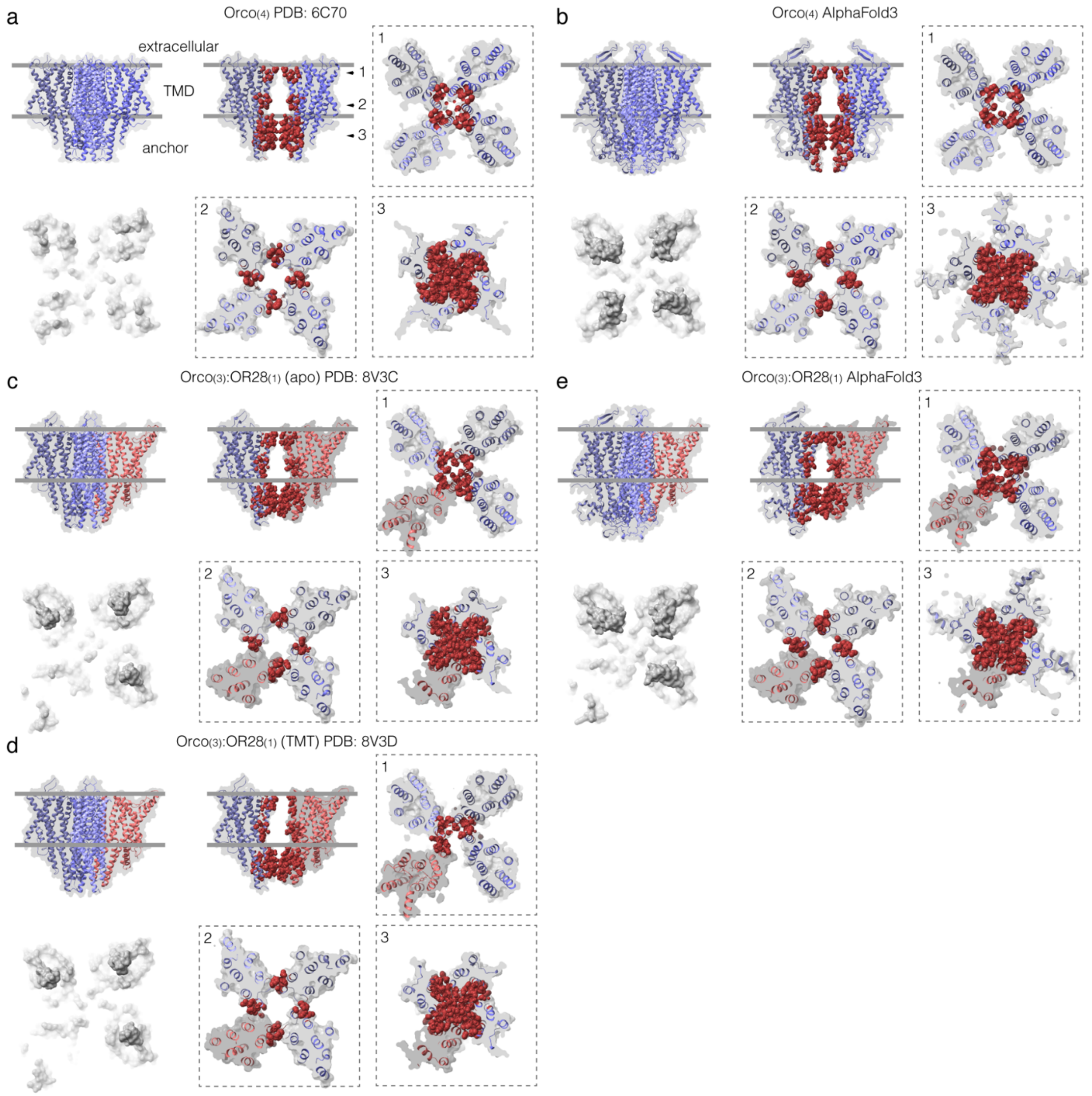


**Supplementary Figure 1| Structures of insect odorant receptors. (a)** and **(b)** $Orco_{(4)}$ homo-tetramer structural models from cryo-EM (a) (PDB: 6C70[1]) and Alphafold3[2] prediction (b). **(c)** to **(e)** $Orco_{(3)}$/$OR28_{(1)}$ hetero-tetramer structural models from cryo-EM in apo (c) (PDB: 8V3C[3]) and ligand-bound (2,4,5-trimethylthiazole, TMT) conditions (d) (PDB: 8V3D[3]), and from Alphafold3 prediction in apo (e). Each panel includes a side view (top left), the extracellular side membrane-protruding surface (bottom left), and OR receptors protomer interface interaction analyses. From 1 to 3: Cross-sections at the extracellular gate (1), transmembrane domain (2), and cytosolic anchor-domain (3). Interface atoms within 5Å of the neighbor Orco/OR28 protomer are shown as red spheres. $Orco_{(4)}$ and $Orco_{(3)}$/$OR28_{(1)}$ share similar structural architecture, encompassing an extracellular domain, a transmembrane domain (TMD), and a cytosolic anchor-domain. The extracellular beta-sheet structure of Orco (which is particularly important for HS-AFM analysis) was not resolved in the $Orco_{(4)}$ cryo-EM structure and partially resolved in the $Orco_{(3)}$/$OR28_{(1)}$ cryo-EM structure. Therefore, Alphafold3-predicted models were provided to supplement the structural analysis. Both $Orco_{(4)}$ and $Orco_{(3)}$/$OR28_{(1)}$ are tethered mainly in the anchor-domain, as highlighted by the large number of interactions at this interface. Protomers are loosely packed in the transmembrane domain, resulting in a unique architecture as compared to canonical tetrameric ion channels.

17

### Supplementary Figure 2

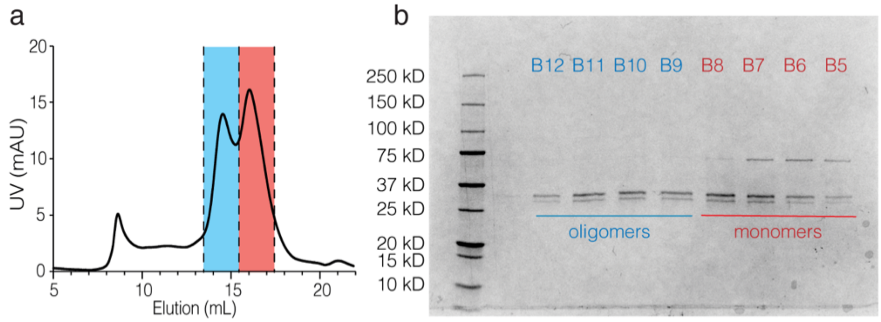


**Supplementary Figure 2| Purification of *Abak*Orco V469A mutant. (a)** Size-exclusion chromatography (SEC) elution profile of the *Abak*Orco Val469Ala (V469A) homo-receptors. The chromatogram highlights two major elution populations. The primary peak (blue-shaded region) corresponds to the assembled oligomeric complex. The secondary peak (red-shaded region) presumably corresponds to the monomeric species. **(b)** SDS-PAGE analysis of fractions collected pose-SEC purification. Fractions B12-B9, corresponding to the primary peak in (a), contain the purified oligomeric complex, whereas fractions B8-B5 represent the presumable monomeric population (secondary peak).

### Supplementary Figure 3

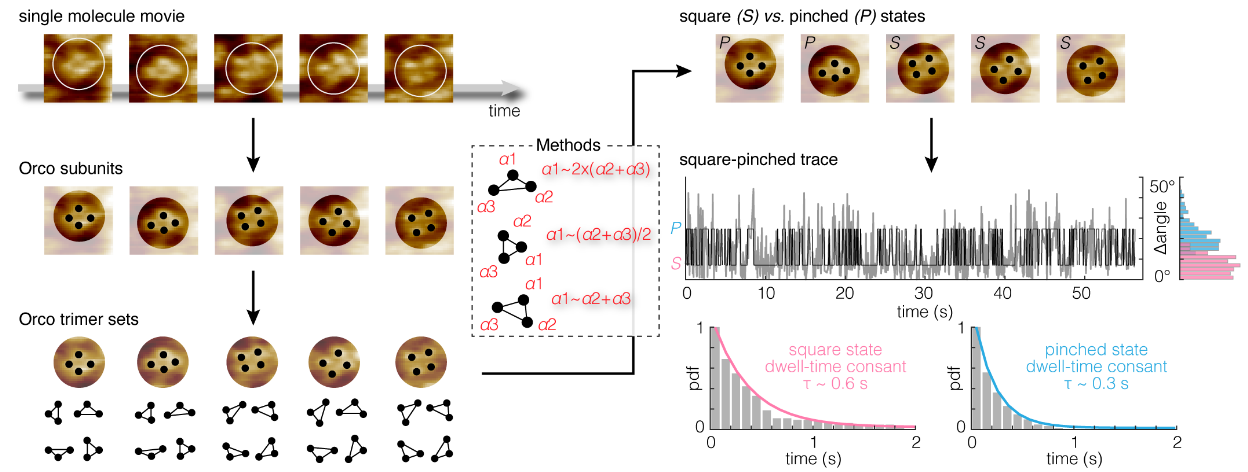


**Supplementary Figure 3| Computational workflow of square and pinched conformational state classification and analysis.** Single molecules were tracked from the HS-AFM movie, where Orco membrane-protruding subunits were located on each single particle. The Orco subunits were then allocated into trimer sets. Each $Orco_{(4)}$ homo-tetramer (shown here) has four such trimer subsets, while each $Orco_{(3)}/OR_{(1)}$ hetero-tetramer has only one. The inter-subunit angles of the trimer sets were used as a criteria to classify single-particles into square and pinched conformational states (**Methods**). Then, for each single molecule, a square-pinched two-state time-trace was constructed, from which the dwell-time contants of square and pinched states were determined by fitting a single exponential to the dwell-time cumulative probability distribution (pdf) histograms.

#### Supplementary Figure 4

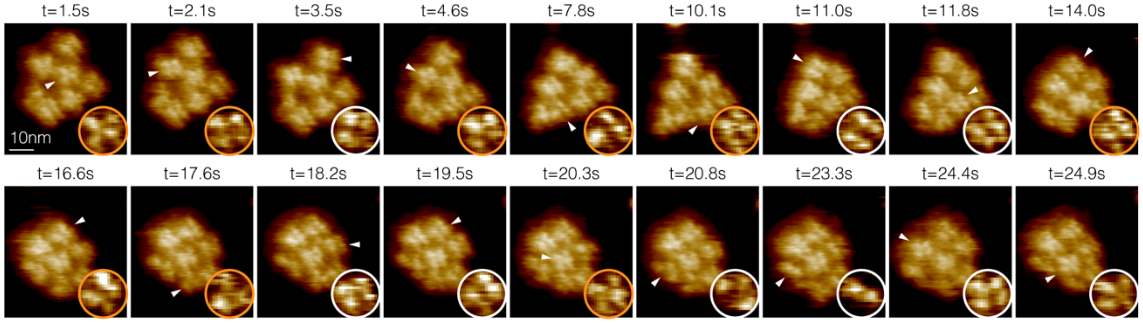


**Supplementary Figure 4| Assembly and conformational dynamics of Orco homo-oligomers with agonist VUAA1.** HS-AFM imaging of membrane-reconstituted Orco homo-oligomeric receptors in the presence of its agonist VUAA1 (0.1 mM, **Supplementary Movie 4**). Imaging parameters: 100 ms/frame, 0.5 nm/pixel.

### Supplementary Figure 5

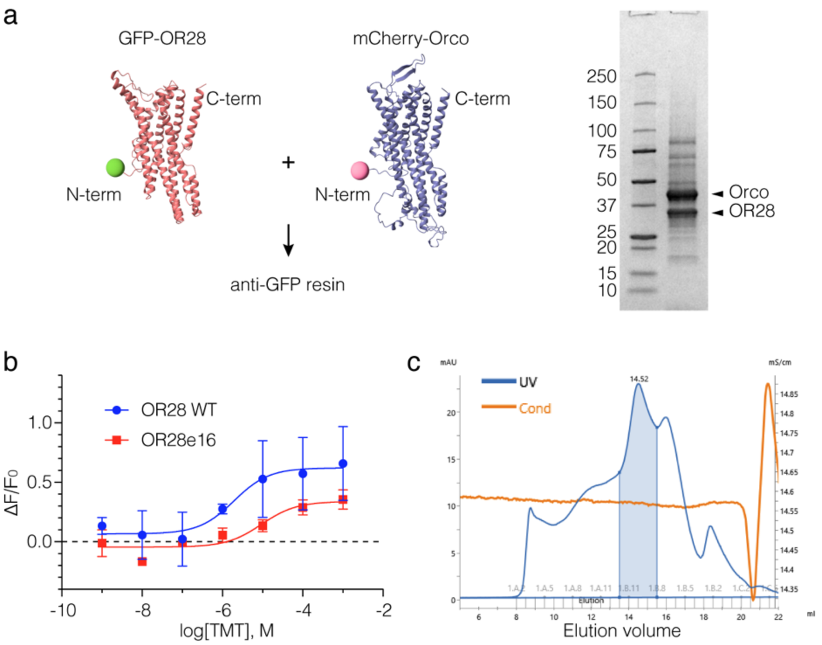


**Supplementary Figure 5| Purification and charaterization of the Orco/OR28 heteromeric receptor samples. (a)** WT Orco/OR28 and engineered Orco/OR28e16 purifications employ an anti-GFP affinity purification targeting specifically GFP-OR28 subunits. **(b)** GCaMP assay showing the 2,4,5-trimethylthiazole (TMT) dose-response curves of WT and engineered OR28 in the presence of Orco. Error bars represent s.e.m from technical replicates with n = 2**. (c)** Size-exclusion chromatography (Superose 6 10/300) profile of the Orco/OR28e16 complex purification. Fractions of the shaded region were pooled for HS-AFM studies.

### Supplementary Figure 6

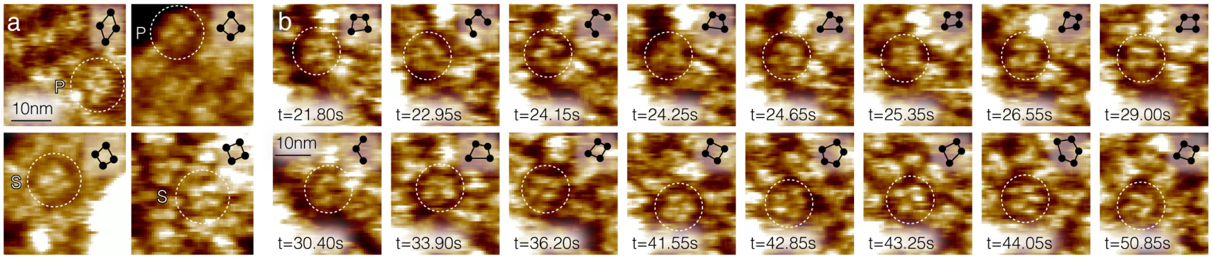


**Supplementary Figure 6| Higher-order Orco oligomers (>3 Orco subunits) in the Orco/OR28 sample. (a)** $Orco_{(4)}$ in pinched (top), and square (bottom) states. **(b)** HS-AFM image series of active Orco homo-oligomer assembly rearrangements in the Orco/OR28 sample (**Supplementary Movie 18**). The Orco homo-oligomer was visualized to undergo active transitions among the $Orco_{(3)}$ (t = 30.40 s), $Orco_{(4)}$ (t = 21.80 s to t = 29.00 s, t = 33.90 s to t = 41.55 s, t = 43.25 s, and t = 50.85 s), and $Orco_{(5)}$ (t = 42.85 s and t = 44.05 s) states. Imaging parameters: 50 ms/frame, 0.67 nm/pixel

28 **Supplementary Figure 7**

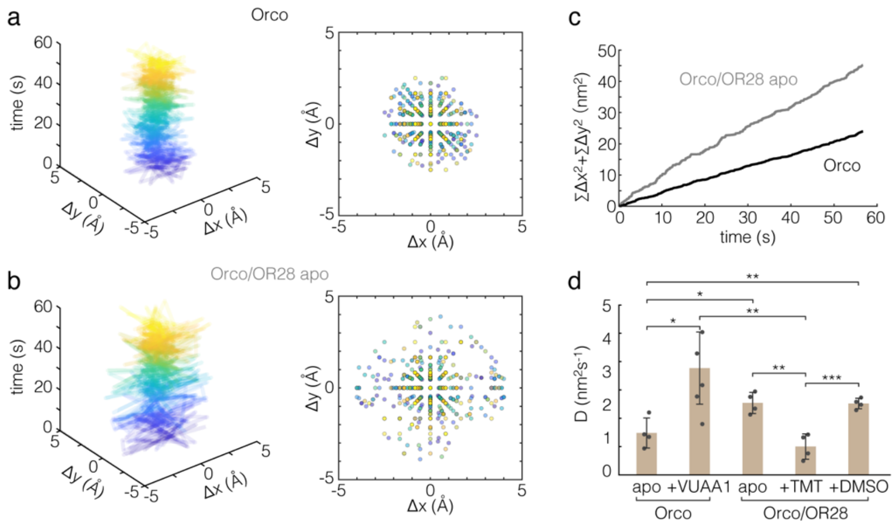


**Supplementary Figure 7| Mobility of individual ORs in clusters. (a)** and **(b)** Drift-corrected time-evolved stepwise (delta) displacement of representative molecules within the protein clusters: an Orco homo-oligomer (top) and an Orco/OR28 hetero-oligomer in apo condition (bottom). Left: Time-evolved delta displacement trace. Right: Distribution of delta displacement. Delta displacements characterize the changes in the *x*- and *y*-dimensions from frame *n* to frame *n*+1 of individual molecules. **(c)** Cumulative delta displacements of representative molecules. **(d)** Diffusion coefficients (mean±s.d.) as a characteristic value for the mobility of individual molecules in the clusters (**Methods**). Two-tailed student *t*-test: *n.s.* not significant (not labelled), $^{*}P<0.05$, $^{**}P<0.01$, $^{***}P<0.001$.

29

### Supplementary Figure 8

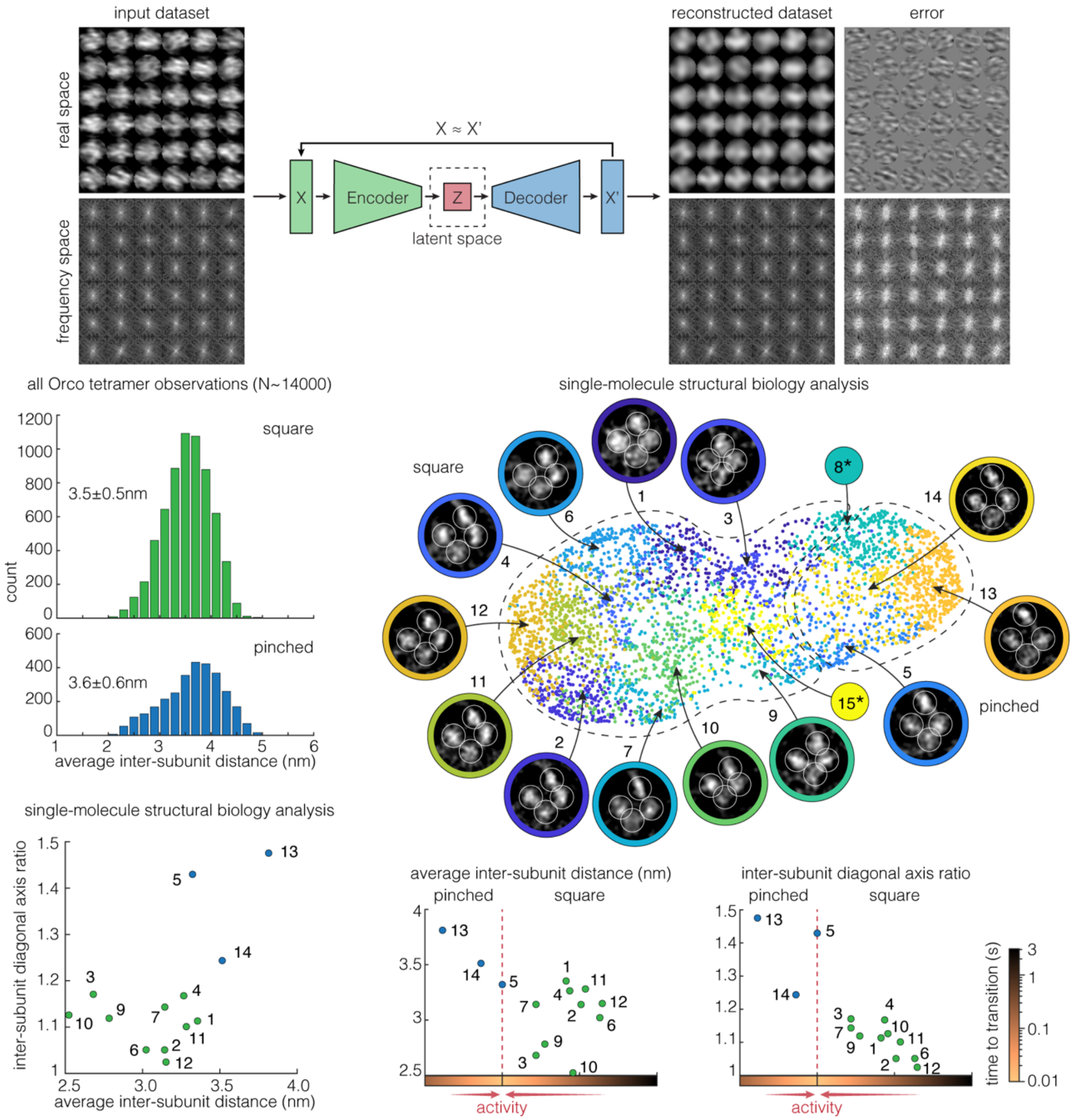


**Supplementary Figure 8| HS-AFM single-particle conformational clustering with an auto-encoder (AE) network.** AE networks are designed to project high-dimensional data (*X*) onto a low-dimensional 'latent space' (*Z*), via an 'encoder', and then reconstruct the original input from this compressed representation (*X'*) with minimal loss (*X~X'*), via a 'decoder'. A well-trained AE network captures the essential features necessary to distinguish between images in its latent space, clustering similar images together while filtering out noise and irrelevant information in an unsupervised manner. Leveraging its power, we implemented an AE-based neural network for HS-AFM single-particle image classification (**Methods**). To discourage biased or irrelevant-feature learning, the input dataset (3,932 input image stack) was augmented with random rotations and translations during training. As a result, the reconstructed dataset effectively captured essential single-particle structural/image features, while the error dataset retained irrelavant and biased information (*e.g.*, scan lines, anisotropic particle orientations, and alignment errors). The resulting latent space was used for further single particle classification (see **Figure 6c-e**, **Methods**). The 2D U-map representation of the latent space from the autoencoder-based (AE) neural network (see **Figure 6b,c**), where a total of 15 conformational classes were revealed and their localization AFM (LAFM) maps calculated (**Methods**). Particles in clusters 8 and 15 (asterisks) gave LAFM maps of poor quality and hence were omitted from substate assignment. Instead, particles in these clusters were classified to either square or pinched states based on their nearest neighbors in the time trace.

32 **Supplementary Figure 9**

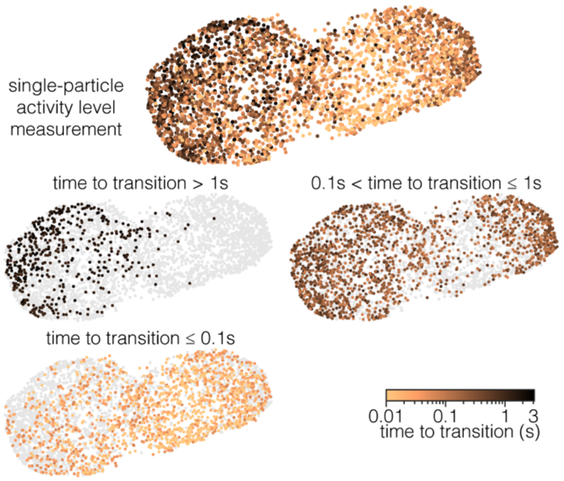


**Supplementary Figure 9| HS-AFM single-particle conformational activity.** 'Time-to-transition' is defined as the shortest temporal distance associated with each particle to a square-pinched state transition in the two-state conformation-time trace (see **Figure 6f**). This metric evaluates the the activity of individual particles. The distribution of the single particles with varying activity levels (false color scale) is displayed on the AE latent space constructed solely on their image/structural features (see **Figure 6c-e**, **Extended Data Figure 8**). Thus, this map reflects the conformation-activity relationship of the molecule.

33

### Supplementary Figure 10

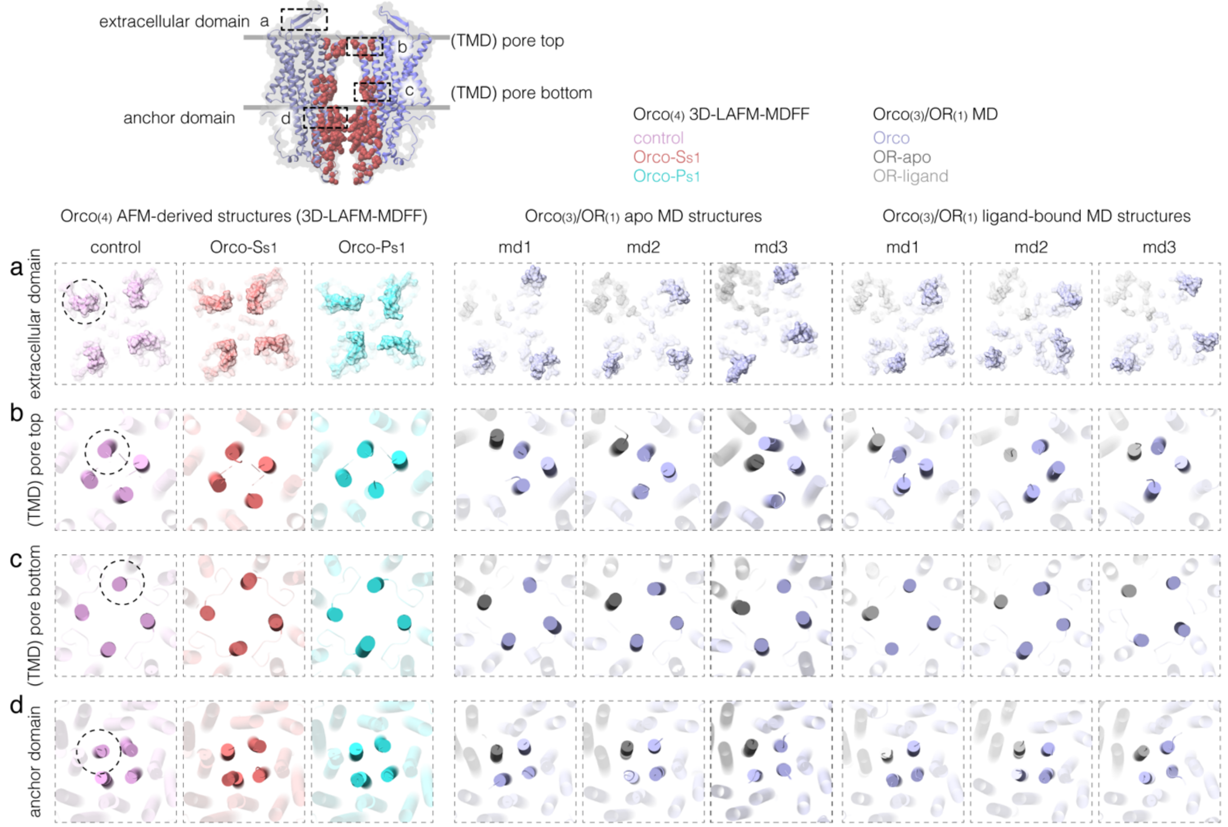


**Supplementary Figure 10| Structural comparisons of MD-derived Orco/OR tetramers.** Comparisons between AFM-derived (3D-LAFM-MDFF, see **Methods**) structures of $Orco_{(4)}$ in apo (Left) and MD structures[4] of $Orco_{(3)}/OR_{(1)}$ in apo (Middle) and ligand-bound condition (Right). **(a)** to **(d)** Structural domains compared encompass the extracellular domain (a), the central pore near the extracellular side (b) and the cytosolic side (c) within the transmembrane domain (TMD), and the anchor-domain (d). For AFM-derived structures, 3D-LAFM-MDFF force fields applied include *Orco-Ss1* (stable square, megenta), *Orco-Ps1* (stable pinched, cyan), and control (no force field, red). For MD structures: blue: Orco subunit, dark gray: OR subunit in apo, light gray: OR subunit with ligand (three independent replicates are shown for each condition).

### Supplementary Tables

**Supplementary Table 1**

| Orco WT apo | | | | |
|---|---|---|---|---|
| assembly | count | lifetime (s) | % by particle-count | % by particle-lifetime |
| $Orco_{(5)}$ | 7 | 138.9 | 8 | 8 |
| $Orco_{(4)}$ | 58 | 1269.5 | 65 | 76 |
| $Orco_{(3)}$ | 22 | 236.95 | 25 | 14 |
| $\leq Orco_{(2)}$ | 2 | 16.5 | 2 | 1.0 |
| Total | 89 | 1661.85 | | |
| **Orco WT VUAA1** | | | | |
| assembly | count | lifetime (s) | % by particle-count | % by particle-lifetime |
| $Orco_{(5)}$ | 0 | 0 | 0 | 0 |
| $Orco_{(4)}$ | 25 | 907.6 | 68 | 72 |
| $Orco_{(3)}$ | 11 | 342.3 | 30 | 27 |
| $\leq Orco_{(2)}$ | 1 | 11.8 | 3 | 1 |
| Total | 37 | 1261.7 | | |
| **Orco WT apo +VUAA1** | | | | |
| assembly | count | lifetime (s) | % by particle-count | % by particle-lifetime |
| $Orco_{(5)}$ | 7 | 138.9 | 6 | 5 |
| $Orco_{(4)}$ | 83 | 2177.1 | 66 | 75 |
| $Orco_{(3)}$ | 33 | 579.25 | 26 | 20 |
| $\leq Orco_{(2)}$ | 3 | 28.3 | 2 | 1 |
| Total | 126 | 2923.55 | | |

**Supplementary Table 1| Population distribution of Orco WT homo-oligomers.** Canoical $Orco_{(4)}$ homo-tetrameric receptor is highlighted

39 **Supplementary Table 2**

| Orco V469A apo | | | | |
|---|---|---|---|---|
| assembly | count | lifetime (s) | % by particle-count | % by particle-lifetime |
| ≥$Orco_{(6)}$ | 4 | 576.9 | 7 | 5 |
| $Orco_{(5)}$ | 14 | 556.7 | 16 | 19 |
| $Orco_{(4)}$ | 20 | 656.2 | 28 | 27 |
| $Orco_{(3)}$ | 21 | 383.5 | 24 | 28 |
| ≤$Orco_{(2)}$ | 16 | 167.8 | 25 | 21 |
| Total | 89 | 2341.1 | | |
| **Orco V469A VUAA1** | | | | |
| assembly | count | lifetime (s) | % by particle-count | % by particle-lifetime |
| ≥$Orco_{(6)}$ | 7 | 174.6 | 7 | 7 |
| $Orco_{(5)}$ | 43 | 1206.95 | 48 | 45 |
| $Orco_{(4)}$ | 20 | 534.6 | 21 | 21 |
| $Orco_{(3)}$ | 20 | 453.7 | 18 | 21 |
| ≤$Orco_{(2)}$ | 5 | 167.55 | 7 | 5 |
| Total | 95 | 1261.7 | | |

**Supplementary Table 2| Population distribution of Orco V469A homo-oligomers.** Canoical $Orco_{(4)}$ homo-tetrameric receptor is highlighted

40

**Supplementary Table 3**

| Orco/OR28e16 apo | | | | |
|---|---|---|---|---|
| assembly | count | lifetime (s) | % by particle-count | % by particle-lifetime |
| $Orco_{(6)}$ | 5 | 121 | 6 | 5 |
| $Orco_{(5)}$ | 4 | 117.75 | 5 | 5 |
| $Orco_{(4)}$ | 6 | 143.4 | 7 | 6 |
| $Orco_{(3)}$ | 8 | 228.55 | 10 | 10 |
| $\leq Orco_{(2)}$ | 8 | 187.85 | 10 | 8 |
| $Orco_{(4)}/OR_{(1)}$ | 4 | 111.3 | 5 | 5 |
| $Orco_{(3)}/OR_{(1)}$ | 47 | 1409.3 | 56 | 60 |
| $Orco_{(2)}/OR_{(1)}$ | 2 | 47.2 | 2 | 2 |
| $Orco_{(2)}/OR_{(2)}$ | 0 | 0 | 0 | 0 |
| Total | 84 | 2366.35 | | |

| Orco/OR28e16 +TMT | | | | |
|---|---|---|---|---|
| assembly | count | lifetime (s) | % by particle-count | % by particle-lifetime |
| $Orco_{(6)}$ | 1 | 53.8 | 1 | 2 |
| $Orco_{(5)}$ | 4 | 165.3 | 4 | 5 |
| $Orco_{(4)}$ | 5 | 89.8 | 5 | 3 |
| $Orco_{(3)}$ | 17 | 431.2 | 16 | 12 |
| $\leq Orco_{(2)}$ | 9 | 178.4 | 9 | 5 |
| $Orco_{(4)}/OR_{(1)}$ | 2 | 87.6 | 2 | 3 |
| $Orco_{(3)}/OR_{(1)}$ | 63 | 2265.1 | 60 | 65 |
| $Orco_{(2)}/OR_{(1)}$ | 3 | 190.8 | 3 | 6 |
| $Orco_{(2)}/OR_{(2)}$ | 1 | 39.1 | 1 | 1 |
| Total | 105 | 3501.1 | | |

| Orco/OR28e16 apo+TMT | | | | |
|---|---|---|---|---|
| assembly | count | lifetime (s) | % by particle-count | % by particle-lifetime |
| $Orco_{(6)}$ | 6 | 174.8 | 3 | 3 |
| $Orco_{(5)}$ | 8 | 283.05 | 4 | 5 |
| $Orco_{(4)}$ | 11 | 233.2 | 6 | 4 |
| $Orco_{(3)}$ | 25 | 659.75 | 13 | 11 |
| $\leq Orco_{(2)}$ | 17 | 366.25 | 9 | 6 |
| $Orco_{(4)}/OR_{(1)}$ | 6 | 198.9 | 3 | 3 |
| $Orco_{(3)}/OR_{(1)}$ | 110 | 3674.4 | 58 | 63 |
| $Orco_{(2)}/OR_{(1)}$ | 5 | 238 | 3 | 4 |
| $Orco_{(2)}/OR_{(2)}$ | 1 | 39.1 | 1 | 1 |
| Total | 189 | 5867.45 | | |

**Supplementary Table 3| Population distribution of Orco/OR28e16 oligomers.** Canoical $Orco_{(3)}/OR_{(1)}$ hetero-tetrameric receptor is highlighted
